# Single-cell Sequencing Reveals Brain/Spinal Cord Oligodendrocyte Precursor Heterogeneity and Requirement for mTOR in Cholesterol Biosynthesis and Myelin Maintenance

**DOI:** 10.1101/2020.10.22.349209

**Authors:** Luipa Khandker, Marisa A. Jeffries, Yun-Juan Chang, Marie L. Mather, Jennifer N. Bourne, Azadeh K. Tafreshi, Isis M. Ornelas, Ozlem Bozdagi-Gunal, Wendy B. Macklin, Teresa L. Wood

**Affiliations:** Department of Pharmacology, Physiology & Neuroscience, New Jersey Medical School, Rutgers University, Newark, New Jersey; Office of Advance Research Computing, Rutgers University, Piscataway, New Jersey; Department of Cell and Developmental Biology, University of Colorado School of Medicine, Aurora, Colorado; Department of Psychiatry, New Jersey Medical School, Rutgers University, Newark, New Jersey

## Abstract

Brain and spinal cord oligodendroglia have distinct functional characteristics, and cell autonomous loss of individual genes can result in different regional phenotypes. However, sequencing studies to date have not revealed distinctions between brain and spinal cord oligodendroglia. Using single-cell analysis of oligodendroglia during myelination, we demonstrate that brain and spinal cord precursors are transcriptionally distinct, defined predominantly by cholesterol biosynthesis. We further identify mechanistic target of rapamycin (mTOR) as a major regulator promoting cholesterol biosynthesis in oligodendroglia. Oligodendroglial-specific loss of mTOR compromises cholesterol biosynthesis in both the brain and spinal cord. Importantly, mTOR loss has a greater impact on cholesterol biosynthesis in spinal cord oligodendroglia that corresponds with more pronounced developmental deficits. However, loss of mTOR in brain oligodendroglia ultimately results in oligodendrocyte death, spontaneous demyelination, and impaired axonal function, demonstrating that mTOR is required for myelin maintenance in the adult brain.

## Main

Oligodendrocytes are the myelin-producing cells of the central nervous system (CNS). There are multiple lines of evidence that oligodendrocytes and oligodendrocyte precursor cells (OPCs) from different regions of the CNS have distinct origins and are functionally dissimilar, which has implications for myelination and regenerative capacity^1,2^. For example, OPCs from the cortex and spinal cord produce myelin sheaths of different lengths when cultured in similar conditions^1^, and transplanted OPCs from white matter differentiate faster than OPCs from gray matter^3^. However, the molecular and cellular mechanisms underlying these phenotypic differences are not known. Recent advances in single-cell transcriptomics have revealed heterogeneous oligodendroglial populations, yet no clear distinction has been shown between brain and spinal cord oligodendroglia in these reports^4,5^.

In addition to distinct developmental origins and cell intrinsic functional differences in oligodendroglia in brain and spinal cord, lineage-specific gene deletions also can vary in phenotype between the two CNS regions. The mechanistic target of rapamycin (mTOR) signaling pathway regulates developmental myelination, but the oligodendroglial specific deletion of *mTOR,* or the mTORC1 binding partner *Raptor,* results in hypomyelination in the spinal cord but not the brain^6–8^. Despite some disruption of myelin gene expression, myelin ultrastructure forms normally in the callosum when *mTOR* or *Raptor* are ablated from developing oligodendroglia^6,8^.

Here we define cellular mechanisms which establish distinctions between brain and spinal cord oligodendroglia and demonstrate that during the crucial stage of developmental myelin initiation, oligodendrocyte precursor populations in spinal cord exhibit higher levels of cholesterol biosynthesis than the equivalent stage precursors in the brain. We also demonstrate that mTOR promotes cholesterol biosynthesis in oligodendroglia. These data support the conclusion that there is a differential requirement for cell-autonomous cholesterol biosynthesis in the brain and spinal cord, and that decreased dependence on cholesterol biosynthesis allows brain oligodendrocytes to myelinate when mTOR is deleted. However, over time, mTOR loss in the corpus callosum results in demyelination and axonal dysfunction, demonstrating that mTOR is required for myelin maintenance in the adult brain.

## Results

### Single-cell sequencing of O4+ oligodendroglia to analyze regional heterogeneity and effects of mTOR deletion

To investigate oligodendroglial diversity during developmental myelination, we isolated the dynamic population of cells identified by the O4 antigen from mouse brains and spinal cords during active developmental myelination. In the developing rodent CNS, O4 is a cell surface sulfatide expressed on late stage OPCs after the early markers platelet-derived growth factor receptor alpha (PDGFRa) and neural/glial antigen 2 (NG2) are mostly downregulated that continues to be present on pre-myelinating as well as actively myelinating oligodendrocytes^9–11^. In order to analyze stage-matched oligodendroglia in the two CNS regions, we magnetically isolated O4+ cells from spinal cord at postnatal day 10 (P10) and brain at P14 since the spinal cord myelinates earlier than the brain^12^. These timepoints correspond to similar developmental phases in the two regions when the most rapid phase of myelination begins^13^. In addition, we analyzed the proportion of cells in the P10 spinal cord and P14 brain that have matured to an O4+ stage (Supplementary Fig 1a, b) and confirmed that P10 spinal cord and P14 brain oligodendroglia have similar percentage of O4+ cells (64.8% of P10 spinal cord cells are O4+ and 60.3% of P14 brain cells are O4+). We used single-cell sequencing of the isolated O4+ oligodendroglia during developmental myelination to 1) reveal transcriptional changes underlying heterogeneity between brain and spinal cord both in normal CNS and after oligodendroglia-specific deletion of mTOR, and 2) to define the pathways which require mTOR for cellular functions necessary for terminal maturation and myelination.

We utilized 10x Genomics platform for single-cell RNA-Seq (Supplementary Fig 2a). After filtering for cell type and quality (see methods), 8,208 control O4+ cells (3,605 from P10 spinal cord, 4,603 from P14 brain) and 9,120 mTOR cKO *(mTOR^fl/fl^; Cnp-Cre* described in methods) O4+ cells (5,233 from P10 spinal cord, 3,887 from P14 brain) were plotted using Uniform Manifold Approximation and Projection (UMAP) for visualization of clustered oligodendroglial populations. Control brain and control spinal cord O4+ cells were integrated together to analyze regional heterogeneity on one UMAP plot, and all four samples (control brain and spinal cord, mTOR cKO brain and spinal cord) were plotted integrated on a second UMAP plot to analyze effects of mTOR loss.

### Regional heterogeneity: During development, brain and spinal cord precursors are transcriptionally distinct

Unsupervised clustering of control O4+ cells revealed 9 distinct populations in developing brain and spinal cord (Fig. 1a). Cell populations were manually identified (Supplementary Fig. 3) using the top 5 differentially expressed genes in each cluster and matched to two reference databases^5,14^. We determined that the clusters form a differentiation continuum of two OPC populations (clusters 7, 8), two committed OPC (COP) populations (clusters 6, 0), a population transitioning from COP to newly formed oligodendrocytes (NFOL) (cluster 1), an NFOL population (cluster 3), a myelin forming oligodendrocyte population (MFOL) (cluster 4), a population transitioning from MFOL to mature oligodendrocyte (MOL) (cluster 5), and finally an MOL population (cluster 2) (Fig 1a, Supplementary Fig. 2c, Supplementary Fig. 3). Our clustering data indicated that the overall distribution of oligodendroglial cell populations was comparable between the isolated brain and spinal cord O4+ cells (Supplementary Fig. 2d). There is a minor shift towards more precursors (OPCs and COPs) in the brain (12.7% difference), equivalent proportions of NFOLs, and MFOLs in the two CNS regions, and a small shift towards more MOLs in the spinal cord (11.2% difference). These data support the conclusion that the two regions are closely stage-matched in terms of oligodendroglial differentiation.

**Fig. 1.**
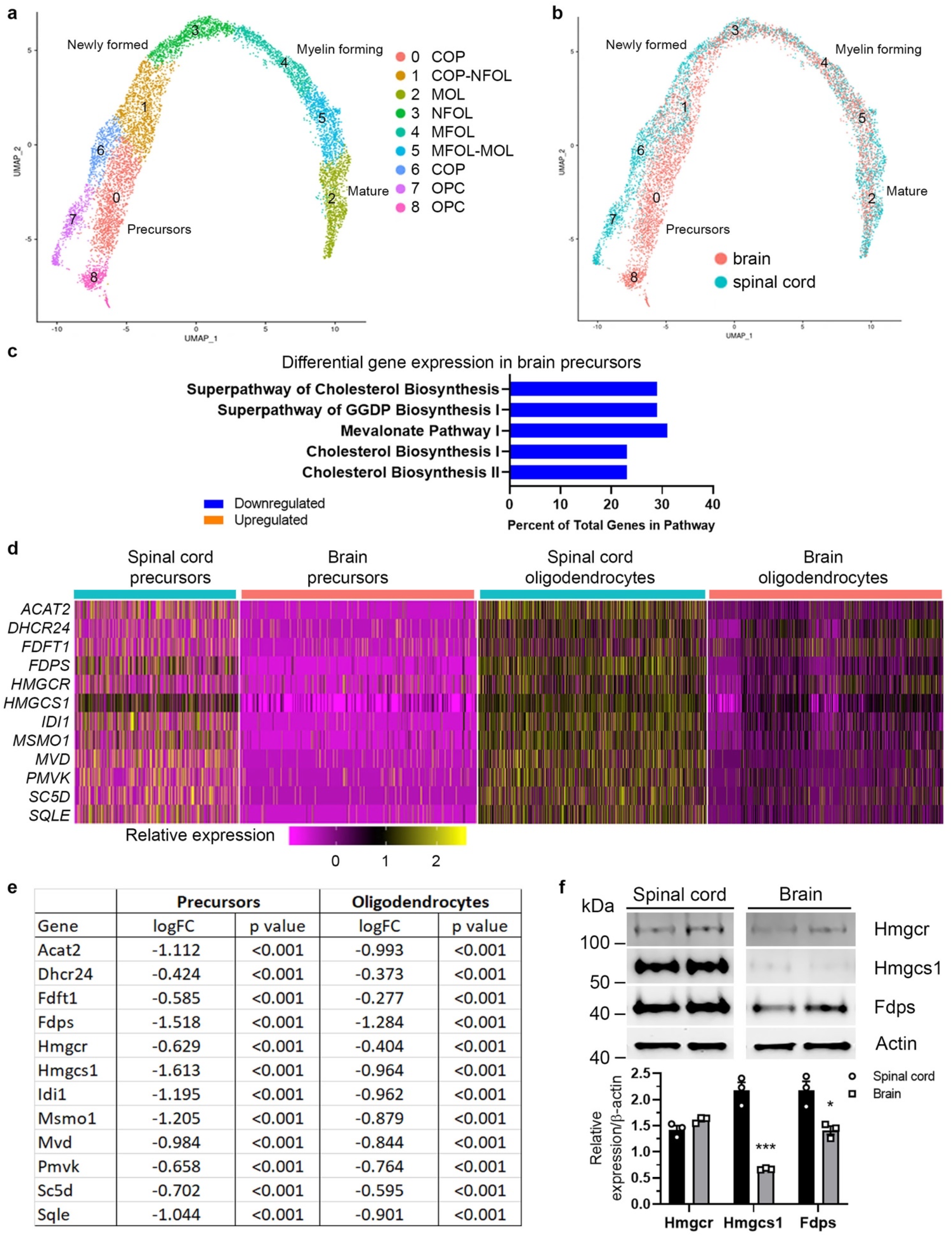
Single-cell sequencing of O4+ oligodendroglia from developing CNS reveals distinct populations in brain and spinal cord; brain precursors have lower expression of cholesterol biosynthesis enzymes compared to spinal cord. **a**, UMAP plot of 8,208 O4+ oligodendroglia from developing brain (P14) and spinal cord (P10). Cells cluster along a differentiation continuum from oligodendrocyte precursor cells (OPC) to committed OPCs (COP), newly formed oligodendrocytes (NFOL), myelin forming oligodendrocytes (MFOL), and finally mature oligodendrocytes (MOL). 3 littermates pooled for brain; 4 littermates pooled for spinal cord. **b**, O4+ cells from brain (4603 cells) and spinal cord (3605 cells) cluster distinctly at precursor stages and converge as they mature. Pink: brain; blue: spinal cord. **c**, Brain precursors (clusters 8 and 0) were compared to spinal cord precursors (clusters 7 and 6), differentially expressed genes were analyzed using IPA, and the top 5 global pathways are shown. Blue: genes with lower expression in brain; orange: genes with higher expression in brain. **d**, Heatmap showing relative single-cell expression of 12 cholesterol biosynthesis genes in brain and spinal cord precursors (clusters 8, 7, 0, 6), and oligodendrocytes (clusters 1, 3, 4, 5, 2). **e**, Differential expression of cholesterol biosynthesis genes extracted from single-cell data and presented as log fold change (logFC) in brain compared to spinal cord. Precursors (clusters 8, 7, 0, 6) and oligodendrocytes (clusters 1, 3, 4, 5, 2) compared separately. **f**, Representative western blots showing expression of the cholesterol biosynthesis enzymes HMGCR, HMGCS1 and FDPS in O4+ cells from P10 spinal cord and P14 brain. Spinal cord and brain samples were run on the same western blot. Data quantified from 3 animals/genotype/CNS region. Values expressed as mean ± SEM *p < 0.05; ***p < 0.001

The MOL cells (cluster 2) expressed *Mal* and *Trf (transferrin)* (Supplementary Fig. 2e), genes whose products are important for myelination^15,16^, as well as *Gsn* (*gelsolin*) and *Stmn1* (*stathmin*) which destabilize the actin cytoskeleton or microtubules, respectively^17,18^. Since actin depolymerization promotes myelin wrapping^19^, we concluded that cluster 2 consisted of oligodendrocytes that were actively wrapping axons. Interestingly, cluster 2 was transcriptionally distinct from the six populations of MOLs previously identified in the adult CNS^5^ and instead shared transcriptomic similarity to multiple MOL clusters (Supplementary Fig. 3b, c), indicating a population of mature oligodendrocytes uniquely present during the myelin initiation phase of development.

Figure 1b identifies cells on the UMAP plot originating from brain or spinal cord. Despite several studies that have shown cell intrinsic regional heterogeneity of oligodendroglia, single cell studies to date have not found clear distinctions between the brain and spinal cord^1,4,5,20^. In contrast, we observed that clusters 7 (OPC) and 6 (COP) were primarily comprised of cells from the spinal cord, whereas clusters 8 (OPC) and 0 (COP) were almost exclusively brain oligodendroglia (Fig. 1b). In cluster 1, a population which represented the next phase of differentiation and was comprised of COPs transitioning to NFOLs, brain and spinal cord cells no longer clustered distinctly, although some separation could still be seen within the population. Upon further maturation, brain and spinal cord oligodendrocytes converged and overlapped completely in clusters 3, 4, 5 and 2. The differentiation trajectory of O4+ clusters in our data was consistent with previous single-cell oligodendroglial studies^5^; however, unlike earlier studies, we identified distinctions between brain and spinal cord precursors. It is likely that stage-matching the two regions and analyzing a dynamic developmental timepoint enabled detection of regional heterogeneity in our study.

To determine transcriptional differences underlying the distinct precursor populations of brain and spinal cord, we compared clusters 8 and 0 (brain OPCs and COPs) to clusters 7 and 6 (spinal cord OPCs and COPs) and analyzed the differentially expressed genes using Ingenuity pathway analysis (IPA). The top 5 pathways representing the most significant differences between brain and spinal cord precursors were associated with cholesterol biosynthesis (Fig. 1c, Supplementary Fig. 4a). Strikingly, all 5 of these differentially expressed pathways had significantly lower expression in the brain.

At the gene level, brain precursors had significantly lower expression of 12 cholesterol biosynthesis genes compared to spinal cord precursors (Fig. 1d, e, Supplementary Fig. 4b). These genes included a number of key regulators such as *Hmgcs1,* which encodes the first enzyme in the cholesterol synthesis pathway and showed the largest expression difference between brain and spinal cord. To determine whether the difference in cholesterol biosynthesis gene expression persisted in more mature cells, brain oligodendrocytes (brain cells in clusters 1,3,4,6 and 2) were compared to spinal cord oligodendrocytes (spinal cord cells in clusters 1,3,4,6 and 2). All of the genes maintained lower expression in brain oligodendrocytes compared to spinal cord oligodendrocytes, although the average log fold change (logFC) was greater between the two precursor populations (Fig. 1d, e).

To determine whether the difference in cholesterol biosynthesis gene expression between brain and spinal cord oligodendroglia reflected differential expression of the corresponding enzymes, we measured protein expression in isolated O4+ cells from P10 spinal cord and P14 brains. Hmgcs1 and Fdps showed the largest differences at the transcript level (Fig. 1e) so were included for protein expression analysis along with HMGCR because it is considered the rate-limiting enzyme of cholesterol biosynthesis. Both HMGCS1 and FDPS had significantly lower expression in brain compared to spinal cord oligodendroglia (Fig. 1f). Although decreased at the RNA level, HMGCR protein expression was not significantly different in oligodendroglia from the two regions (Fig. 1f). To rule out the possibility that the lower expression of HMGCS1 and FDPS is due to a more immature state of the cells in the brain versus spinal cord, we isolated brain O4+ cells at P18 and found that the brain cells continued to have lower expression of cholesterol biosynthesis enzymes even at a later developmental stage (Supplementary Fig. 4c). We concluded that brain and spinal cord OPCs and COPs are transcriptionally distinct, and that the main contributing pathways are those associated with cholesterol biosynthesis.

### Oligodendroglial loss of mTOR results in expansion of a population with dysregulated global pathways

To visualize population shifts caused by deletion of mTOR, we reclustered the cells from control samples with mTOR cKO cells as described above (Fig. 2a; Supplementary Fig. 5a). Each cluster was manually identified using the top 5 differentially expressed genes for each cluster (Supplementary Fig. 5c, d) and referencing published databases^5,14^ as for the previous analyses. Similar to the UMAP of control brain and spinal cord, clusters reflected progression along a differentiation continuum from OPCs to COPs to NFOLs to MFOLs and finally to MOLs (Fig. 2a, Supplementary Fig. 5b). The clear distinctions between brain and spinal cord precursors (Fig. 1b) were masked when all samples are combined; this can be attributed to a greater difference in gene expression between control and mTOR cKO cells than between control brain and control spinal cord.

**Fig. 2.**
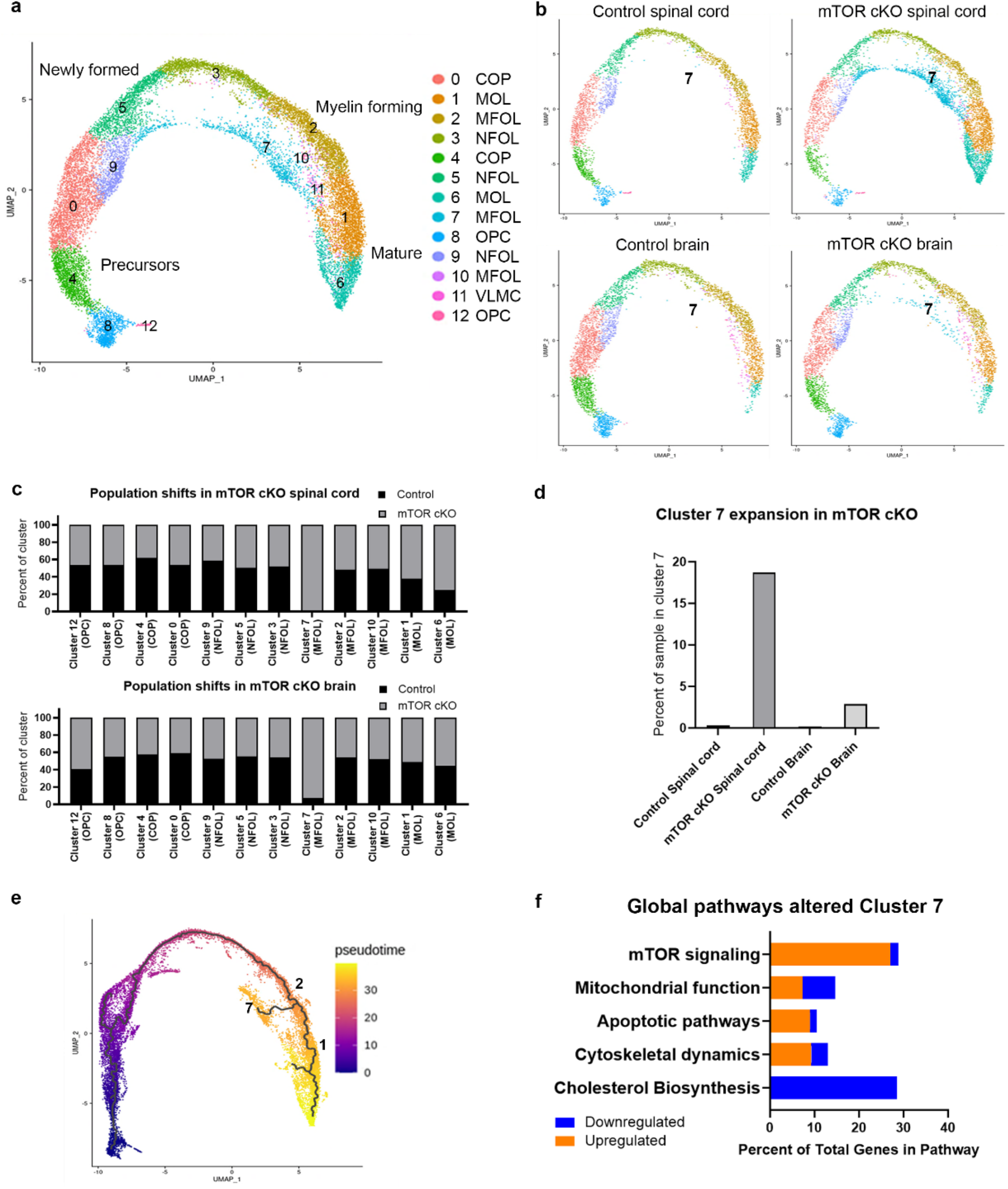
Oligodendroglial loss of mTOR results in expansion of a population with dysregulated global pathways. **a**, Combined UMAP plot of O4+ cells from brain and spinal cord of control and mTOR cKO animals. Cells cluster in a differentiation continuum from OPCs to COPs, NFOLs, MFOLs, and MOLs. Non-oligodendroglial vascular leptomeningeal cell (VLMC) cluster (cluster 11) was excluded from further analyses. OPCs in cluster 12 have a higher expression of cell cycle genes *(Cdca8, Ccnd2, Cks2, Cdkn3, Cdca3, Cdca7, Cdk6),* indicating that they are cycling cells. 8,208 control cells (3,605 from P10 spinal cord, 4,603 from P14 brain); 9,120 mTOR cKO cells (5,233 from P10 spinal cord, 3,887 from P14 brain). 3 littermates/genotype pooled for brain; 4 littermates/genotype pooled for spinal cord. **b**, UMAP plots separated by sample show that cluster 7 is present almost exclusively in mTOR cKO. **c**, Population shifts in mTOR cKO spinal cord and brain. Percent of cells in each cluster that are control or mTOR cKO. **d**, Percentage of each sample present in cluster 7. Control spinal cord: 0.3% (12 cells/3605 total); mTOR cKO spinal cord: 18.7% (980 cells/5233 total); Control brain: 0.2% (9 cells/4603 total); mTOR cKO brain: 2.9% (114 cells/3887 total). **e**, Cells from single-cell analysis are ordered by pseudotime algorithms using Monocle. Trajectory begins at OPCs in purple and progresses to MOLs in yellow. **f**, Top global pathways differentially expressed in cluster 7 compared to cluster 2, analyzed using IPA. Blue: downregulated genes; orange: upregulated genes.

Separating the UMAPs by CNS region allowed us to visualize population shifts resulting from loss of mTOR within each region (Fig. 2b). Although several clusters were altered with deletion of mTOR (Fig. 2c), MFOL cluster 7 showed the largest population shift. In the spinal cord this represented an expansion from 0.3% of control O4+ cells to 18.7% of mTOR cKO O4+ cells in cluster 7; in the brain cluster 7 exhibited a smaller expansion from 0.2% of control to 2.9% of mTOR cKO (Fig. 2d).

We performed pseudotime analysis to identify the origin of cluster 7 cells. Our data indicated that in control oligodendroglia, MFOL cluster 2 progressed to MOL cluster 1 (Supplementary Fig. 6b). In mTOR cKO oligodendroglia, cluster 2 branched to both cluster 1 and cluster 7 (Fig. 2e, Supplementary Fig. 6c). Comparing cluster 7 to its origin, cluster 2, revealed that the top altered pathways fell into one of five categories: mTOR signaling, mitochondrial function, apoptotic pathways, cytoskeletal dynamics, and cholesterol biosynthesis (Fig. 2f). Dysregulation of mTOR signaling was to be expected in a population that was almost exclusively present in mTOR cKO. It was also not surprising that loss of mTOR in differentiating oligodendrocytes impacted genes involved in cytoskeletal reorganization. Our previous studies have defined specific mTOR-regulated cytoskeletal targets in developing oligodendroglia *in vitro* and *in vivo* during both early stages of morphological differentiation and in later stages of myelin wrapping^21,22^. Cytoskeletal dynamics are essential as MFOLs mature to MOLs and must shift from promoting process extension requiring microtubule and actin polymerization to initiating axon wrapping requiring depolymerization^19,23–25^. Our results indicated that cluster 7 cells were unable to undergo the morphological changes necessary for terminal differentiation and initiation of axon wrapping (Supplementary Fig. 6c). Instead of lower expression of *Fyn,* which promotes early morphological differentiation and is downregulated in mature oligodendrocytes^5,14,26^, cells in cluster 7 had higher *Fyn* expression than in cluster 2. In contrast, *Gsn,* which functions in actin-severing, is normally upregulated as an oligodendrocyte progresses from MFOL to MOL^5,14,19^ as seen in cluster 1 (additional supplementary files), but was downregulated in cluster 7. These data are consistent with the impaired initiation of myelination and continued hypomyelination observed in mTOR cKO developing spinal cord^8,21^.

### mTOR loss dysregulates global pathways important for oligodendrocyte biology

Loss of mTOR in developing oligodendroglia caused dysregulation of mitochondrial function, apoptotic pathways and cholesterol biosynthesis based on cluster 7 analyses (Fig. 2f). These pathways are important for normal oligodendrocyte development and function but have not previously been identified as mTOR-regulated in oligodendroglia. Next, we compared all mTOR cKO clusters to all control clusters from spinal cord and identified 61 pathways that were altered; the top 5 pathways are shown in Fig. 3a. Similar to cluster 7, mTOR signaling was dysregulated overall in mTOR cKO spinal cord oligodendroglia. The remaining four pathways, superpathway of cholesterol biosynthesis, superpathway of GGDP biosynthesis I, mevalonate pathway, and cholesterol biosynthesis I, are all associated with cholesterol biosynthesis (Supplementary Fig. 4a).

**Fig. 3.**
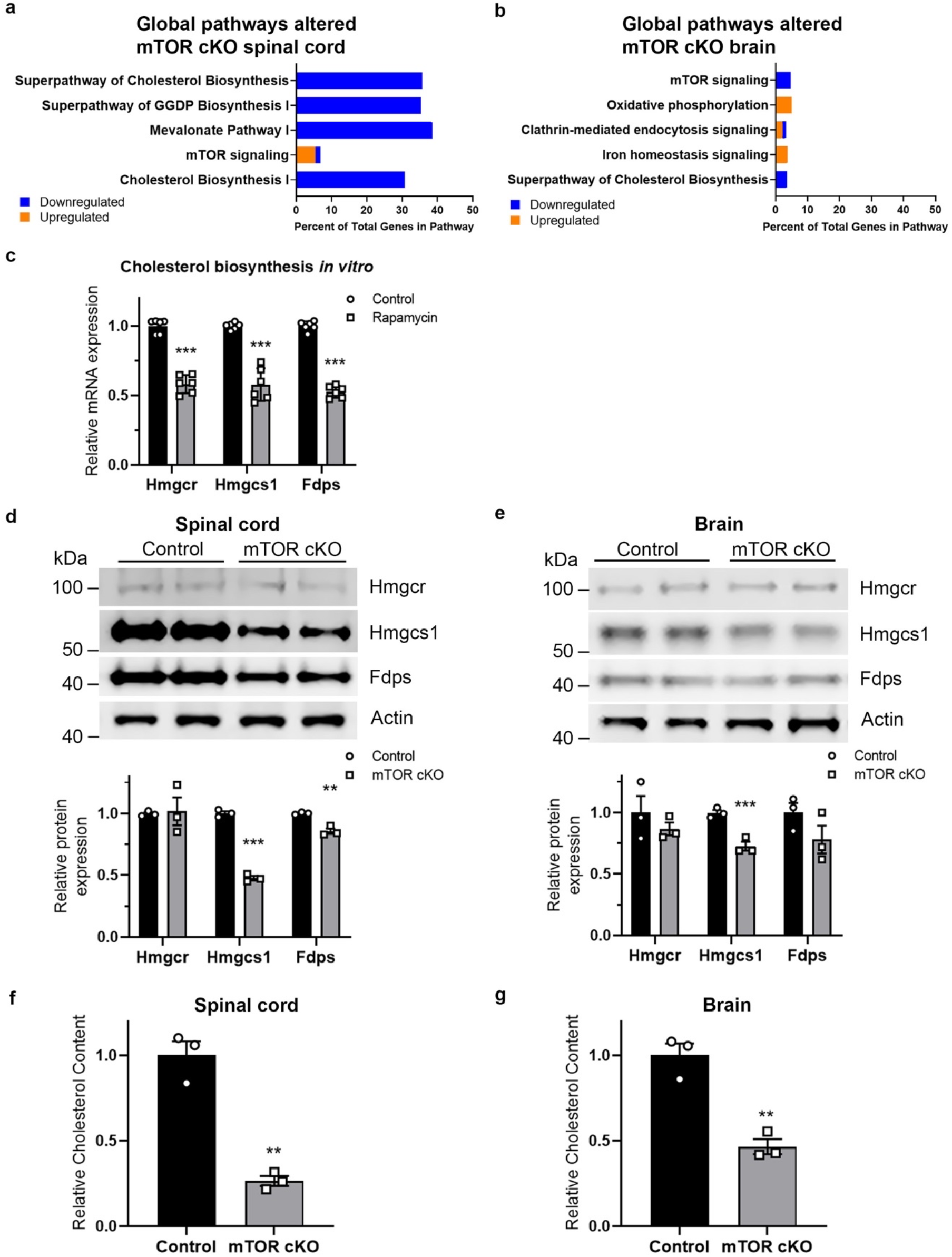
mTOR is necessary for normal cholesterol biosynthesis in oligodendroglia. **a-b**, Top 5 pathways dysregulated in mTOR cKO spinal cord (a) and brain (b). IPA analysis of genes differentially expressed in all mTOR cKO cells compared to all control cells; most significantly altered pathways shown. Blue: downregulated genes; orange: upregulated genes. **c**, Inhibiting mTOR *in vitro* reduces mRNA expression of cholesterol biosynthesis genes. Expression of *Hmgcr, Hmgcs1* and *Fdps* by quantitative real-time PCR in primary rat OPCs differentiated for 3 days in presence or absence of 10nM rapamycin to inhibit mTOR. Data represent 6 replicates over two independent OPC preps. Values expressed as mean ± SD ***p < 0.001. **d-e**, Cholesterol biosynthesis enzymes are reduced in mTOR cKO spinal cord (d) and brain (e), with a greater impact in the spinal cord. Representative western blots showing expression of HMGCR, HMGCS1, and FDPS in control and mTOR cKO spinal cord (d) and brain (e) O4+cells. Expression of cholesterol enzymes was normalized first to β-actin, then to control. Graphs present data from 3 animals/genotype/CNS region. Values expressed as mean ± SEM **p < 0.01; ***p < 0.001. **f-g**, mTOR cKO O4+ cells have lower cholesterol content. Total cholesterol measured in isolated O4+ cells from P10 spinal cords (f) and P14 brains (g) using neutral lipid extraction and enzymatic colorimetric cholesterol assay and normalized to cell number. Graphs present data from 3 animals/genotype/CNS region. Values expressed as mean ± SEM **p < 0.01.

A similar comparison of all mTOR cKO clusters to all control clusters from brain revealed 20 global pathways dysregulated in mTOR cKO brain O4+ cells. As in the spinal cord, mTOR signaling was one of the top 5 pathways (Fig. 3b). The superpathway of cholesterol biosynthesis was also dysregulated in the mTOR cKO brain, but less severely than in the spinal cord with fewer downregulated genes. Genes involved in oxidative phosphorylation, a mechanism for ATP synthesis in the mitochondria, were also dysregulated in mTOR cKO brains.

The remaining top pathways dysregulated in the brain were clathrin-mediated endocytosis signaling, which functions in internalization of molecules from the plasma membrane into intracellular compartments, and iron homeostasis signaling, which regulates iron uptake, storage and export. While both pathways have independent functions important for oligodendrocyte biology, they are also associated with cholesterol homeostasis^27,28^. To determine whether the transcriptomic changes observed in mTOR cKO are due exclusively to the expansion of MFOL cluster 7, we excluded cluster 7 and compared the remaining mTOR cKO cells to control cells and found that global pathways, including mTOR signaling, remained dysregulated (Supplementary Fig. 7a, b). We also considered the possibility that loss of mTOR completely prevents oligodendrocytes from progressing past the MFOL stage, resulting in expansion of cluster 7 and a lack of mTOR cKO MOLs. This could occur if the later MOL cells did not express Cre or did not recombine at the mTOR locus. If so, MOL clusters 1 and 6 would have normal mTOR signaling and cholesterol biosynthesis. To address this, we analyzed genes differentially expressed in mTOR cKO cells exclusively within MOL clusters 1 and 6 (Supplementary Fig. 7c, d). mTOR signaling was dysregulated in mTOR cKO MOLs, indicating that these cells have not escaped mTOR deletion. Moreover, cholesterol biosynthesis was also dysregulated, demonstrating a continued deficit in mTOR cKO MOLs.

Since dysregulation of cholesterol biosynthesis and oxidative phosphorylation were identified in cluster 7 as well as in mTOR cKO overall, we performed functional assays to validate alterations in these pathways with loss of mTOR. Our results indicate that while mTOR regulates oxidative phosphorylation (Supplementary Fig. 8a, b), oligodendrocytes can withstand a small reduction in respiratory capacity without significant effects on myelin gene expression (Supplementary Fig. 8c, d). Moreover, based on the similar oxygen consumption phenotypes in brain and spinal cord, it seems unlikely that this accounts for the greater effect of mTOR loss on spinal cord oligodendroglia in the mTOR cKO.

### mTOR is necessary for normal cholesterol biosynthesis in oligodendroglia

To further investigate mTOR regulation of cholesterol biosynthesis gene expression, we treated differentiating primary OPCs in culture with the mTOR inhibitor rapamycin and measured mRNA expression of *Hmgcr, Hmgcs1,* and *Fdps* (Fig. 3c). Inhibiting mTOR significantly decreased expression of all three cholesterol biosynthesis genes, indicating that mTOR is necessary for their normal expression in oligodendroglia. Next, we assessed protein expression of HMGCR, HMGCS1 and FDPS in O4+ cells isolated from control and mTOR cKO spinal cord (P10) and brain (P14) (Fig. 3d, e). While HMGCR expression was unaffected in mTOR cKO brain and spinal cord, both HMGCS1 and FDPS expression were significantly reduced in mTOR cKO spinal cord. In the brain, mTOR deletion led to a significant reduction in HMGCS1 in oligodendroglia (27% decrease), although reduction was less than in spinal cord (52% decrease). FDPS was unaffected in the developing brain with deletion of mTOR. We concluded that mTOR deletion dysregulates cholesterol biosynthesis enzymes in both brain and spinal cord oligodendroglia but has a greater impact in the spinal cord. To determine whether reduction of cholesterol enzymes lowered cholesterol production, we directly measured cholesterol levels in freshly isolated O4+ oligodendroglia. Total cholesterol was significantly reduced in both mTOR cKO spinal cord and brain O4+ cells (Fig. 3f, g). Consistent with the relative reduction of cholesterol biosynthesis enzymes, loss of mTOR reduced cholesterol content to a greater extent in spinal cord than brain. These findings demonstrate that mTOR promotes expression of cholesterol biosynthesis enzymes, both at the transcript and protein level, and maintains normal cholesterol content.

### Inhibition of cholesterol biosynthesis in primary oligodendroglia decreases cell number and myelin gene expression

Cholesterol is an integral component of the myelin sheath, and decreased cholesterol biosynthesis via pharmacological inhibition or genetic mutation can cause myelin deficits in both the PNS and CNS^29^. To determine whether reduced cholesterol biosynthesis directly affects oligodendrocyte viability or myelin gene expression, we treated differentiating primary OPCs *in vitro* with simvastatin, an inhibitor of HMGCR, the rate-limiting enzyme of the cholesterol biosynthesis pathway. Simvastatin treatment significantly reduced oligodendroglia number in a dose-dependent manner (Fig. 4a). Since most primary oligodendroglia in differentiation conditions are post-mitotic, the reduced cell number from simvastatin treatment was likely due to cell death.

**Fig. 4.**
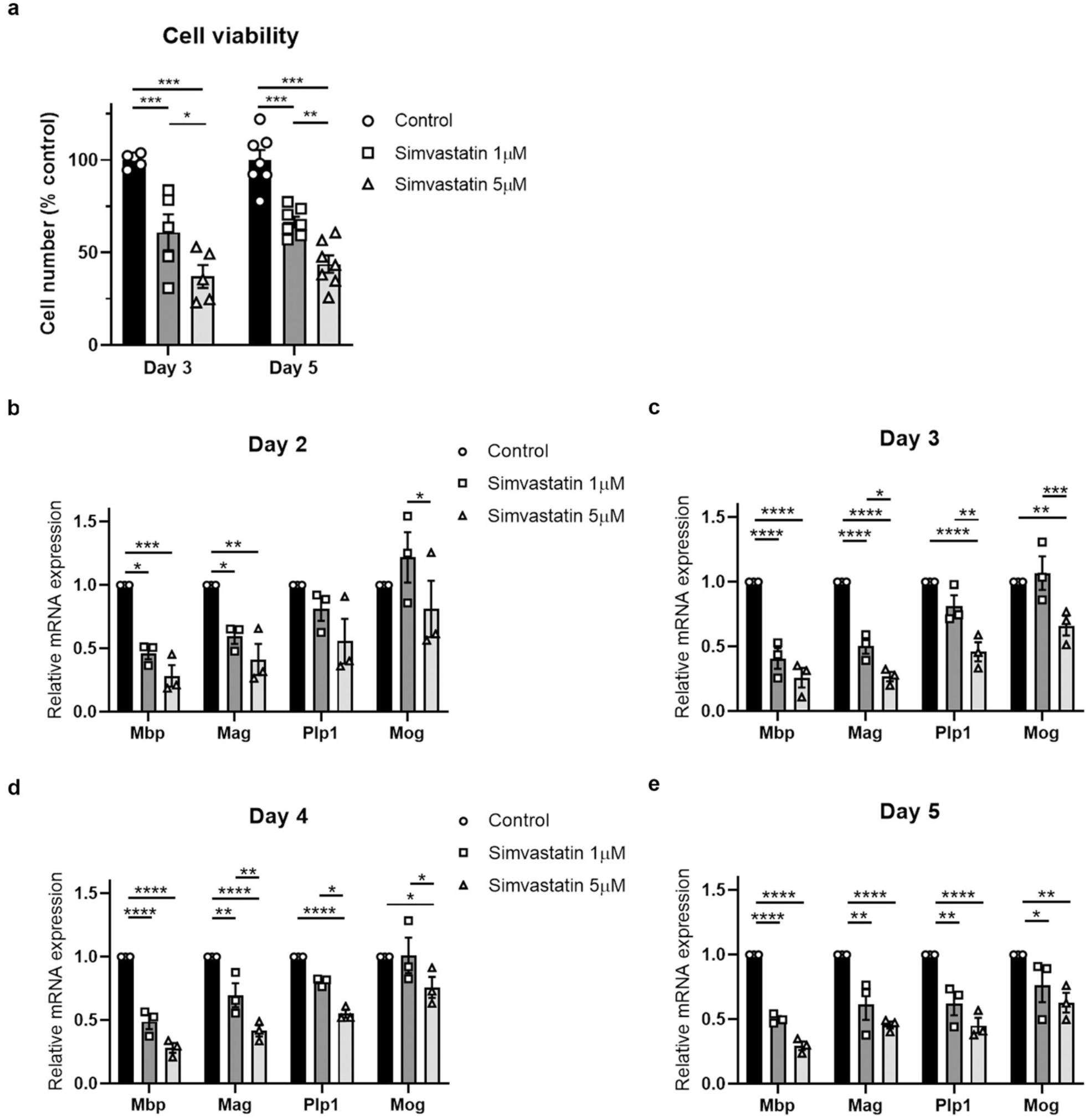
Inhibition of cholesterol biosynthesis in primary oligodendroglia decreases cell number and myelin gene expression. **a**, Inhibition of cholesterol biosynthesis in primary oligodendroglia decreases cell number in dose-dependent manner. Differentiating primary rat OPCs treated with 1μM or 5μM simvastatin (Hmgcr inhibitor) for 3 days and 5 days. Cell viability was measured by MTT assay and normalized to controls. Cells were plated in triplicate and graph represents data from 3 independent experiments. Values expressed as mean ± SEM *p < 0.05; **p < 0.01; ***p < 0.001. **b-e**, Inhibition of cholesterol biosynthesis decreases myelin gene expression in differentiating primary oligodendroglia. Expression of *Mbp* (early marker of oligodendrocyte differentiation), *Mag, Plp1,* and *Mog* (late marker of oligodendrocyte differentiation) measured by quantitative real-time PCR in primary rat OPCs differentiated for 2, 3, 4, and 5 days, treated with 1μM or 5μM simvastatin and compared to control. Expression normalized to *Gapdh* as housekeeping gene, then normalized to control. Data presented from 3 independent experiments. Values expressed as mean ± SEM *p < 0.05; **p < 0.01; ***p < 0.001.

To determine how simvastatin treatment affects myelin gene expression, we measured mRNA expression of *Mbp, Plp1, Mag,* and *Mog* which encode important structural components of myelin in oligodendrocytes differentiating *in vitro* (Fig. 4b-e). Expression of both *Mbp* and *Mag* was significantly reduced in oligodendroglia treated with either 1 μM or 5 μM simvastatin from 2-5 days of differentiation (Fig. 4b-e). *Plp1* and *Mog* expression were reduced only by the higher dose of simvastatin at 3-4 days of differentiation (Fig. 4c, d); by day 5, *Plp1* and *Mog*expression were reduced at both concentrations of simvastatin (Fig. 4e). We conclude that inhibiting cholesterol biosynthesis in primary oligodendroglia with simvastatin reduces myelin gene expression, with higher concentrations and longer treatments having a greater effect. Our findings are consistent with reports that simvastatin administration *in vivo* affects oligodendrocyte numbers and MBP expression during remyelination^30^ and further demonstrate that inhibiting cholesterol biosynthesis has a cell-autonomous effect on oligodendrocyte viability and on expression of essential myelin genes.

### Deficits in cholesterol biosynthesis persist at 8 weeks of age, myelin protein expression is downregulated, and mTOR cKO brain oligodendroglia undergo apoptosis

Our data show that oligodendroglia in the mTOR cKO spinal cord have a greater reduction in cholesterol biosynthesis enzymes and cholesterol content than mTOR cKO brain oligodendroglia. This is consistent with the hypomyelination observed in mTOR cKO spinal cord whereas the brain shows normal developmental myelination^8^. However, it is possible that the lower cholesterol in brain oligodendroglia due to loss of mTOR results in a delayed myelin phenotype.

To first determine whether cholesterol biosynthesis enzymes remained reduced in adult brains, we analyzed microdissected corpus callosum of control and mTOR cKO mice at 8 weeks of age. mTOR cKO corpus callosum exhibited a significant reduction in protein expression of both HMGCS1 and FDPS but not HMGCR (Fig. 5a). We next analyzed expression of MBP and PLP in the corpus callosum at 8 weeks and found that MBP expression was significantly reduced while PLP expression was unchanged (Fig. 5b), revealing effects on myelin proteins similar to those observed *in vitro* (Fig. 4). Additionally, immunostaining for MOG revealed a reduction in area of MOG positivity in the corpus callosum of mTOR cKO animals (Fig. 5c) compared to controls.

**Fig. 5.**
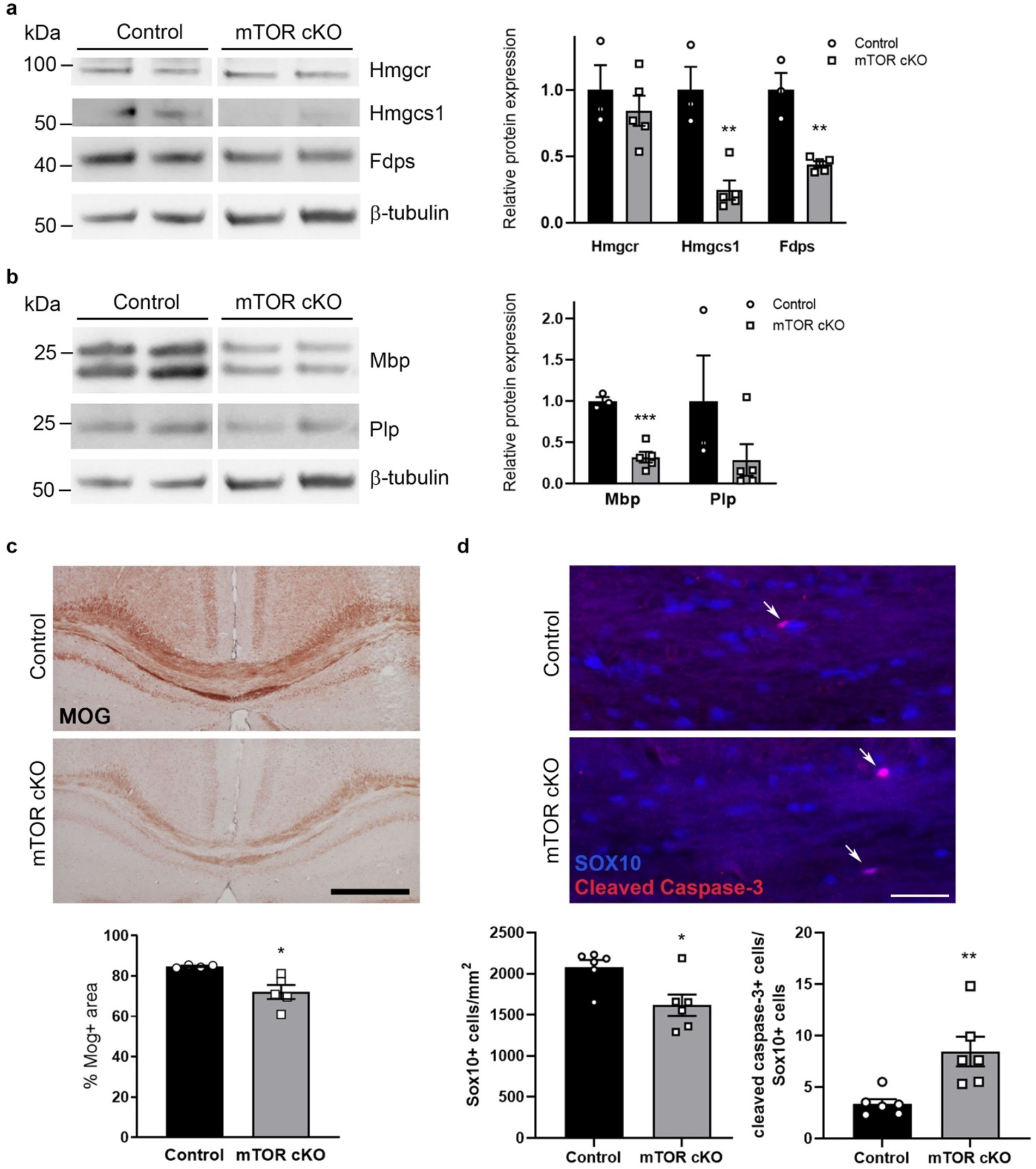
Deficits in cholesterol biosynthesis persist at 8 weeks of age, myelin protein expression is downregulated, and brain oligodendroglia undergo apoptosis. **a**, Cholesterol biosynthesis enzymes are downregulated at 8 weeks in mTOR cKO corpus callosum. Representative western blots showing expression of HMGCR, HMGCS1, and FDPS in control and mTOR cKO microdissected corpus collosum. **b**, Expression of myelin proteins MBP and PLP in mTOR cKO corpus callosum. Representative western blots showing expression of MBP and PLP in control and mTOR cKO microdissected corpus collosum at 8 weeks of age. For **a** and **b**, control and mTOR cKO samples were run on the same western blot. Expression in **a** and **b** was normalized first to β-tubulin, then to control. Graphs for **a** and **b** present data from 3 control and 5 mTOR cKO animals. Values expressed as mean ± SEM **p < 0.01; ***p < 0.001. **c**, MOG expression is dysregulated in mTOR cKO corpus callosum. Representative images of MOG staining shows decreased area of MOG positivity in mTOR cKO corpus callosum compared to control. Scale bar = 500 μm. Graph presents data from 4 control and 5 mTOR cKO animals. All values expressed as mean ± SEM *p < 0.05; **p < 0.01; ***p < 0.001. **d**, Representative images of SOX10 (blue) and cleaved caspase-3 (red) immunostaining from control and mTOR cKO corpus callosum at 7-8 weeks of age, scale bar = 50 μm. Arrows indicate cleaved caspase-3+/ SOX10+ cells. Graphs present data from 6 animals/genotype. All values expressed as mean ± SEM *p < 0.05; **p < 0.01.

Our *in vitro* studies indicate that reduced cholesterol biosynthesis can reduce oligodendrocyte viability (Fig. 4a). Furthermore, apoptotic pathway genes are dysregulated in cluster 7, the population of cells expanded in mTOR cKO (Fig. 2f). Therefore, to determine whether oligodendrocyte survival is affected *in vivo,* we quantified the number of cleaved caspase-3+ cells (a marker of apoptosis) and SOX10+ cells (total oligodendroglial cells) in mTOR cKO and control corpus callosum (Fig. 5d). The number of SOX10+ oligodendroglia was reduced in the corpus collosum of mTOR cKO mice and a corresponding increase in cleaved caspase-3+/SOX10+ cells indicated that mTOR deletion resulted in apoptosis and loss of oligodendroglia in the brain by 8 weeks of age.

### At 12 weeks, mTOR cKO brains exhibit dysregulation of myelin genes and hypomyelination

Our data suggest that although developmental myelination in the mTOR cKO brain is structurally normal by EM at 8 weeks^8^, myelin sheath integrity may be compromised after initial wrapping. Although most developmental myelination is complete by 8 weeks of age in mice, there is continual myelin accumulation between 8 and 12 weeks^31^. Moreover, loss of oligodendrocytes in the adult corpus callosum might further impact myelin integrity. At 12 weeks of age, mTOR cKO mice exhibited significantly reduced MBP and PLP expression compared to controls (Fig. 6a). Immunostaining for MOG also revealed a significant reduction in area of MOG positivity in mTOR cKO (Fig. 6b). These data suggested a progressive myelin deficit from 8 to 12 weeks of age. Analyzing corpus callosal myelination at 12 weeks by EM (Fig. 6c-f), we found fewer myelinated axons (Fig. 6d) and thinner myelin by g-ratio analysis (Fig. 6e) in mTOR cKO. Both linear regression analysis (Fig. 6e) as well as g-ratio distribution analysis (Fig. 6f) revealed shifts toward higher g-ratio values. Taken together with the increased apoptosis at 8 weeks, our data indicate that mTOR deletion from the oligodendrocyte lineage results in oligodendrocyte death and demyelination in the adult brain.

**Fig. 6.**
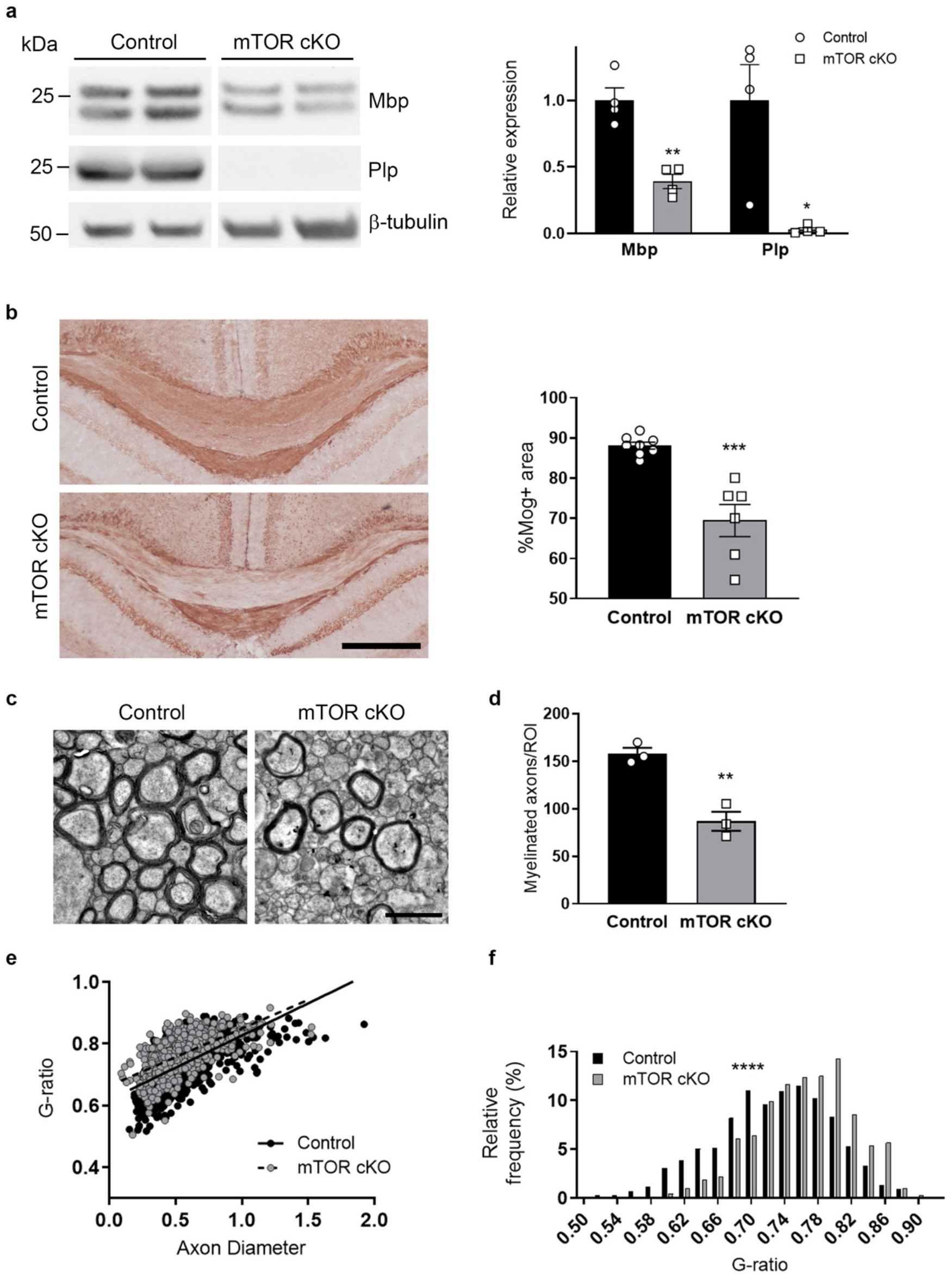
At 12 weeks, mTOR cKO brains exhibit dysregulation of myelin genes and hypomyelination. **a**, Expression of myelin proteins MBP and PLP in mTOR cKO corpus callosum. Representative western blots showing decreased expression of MBP and PLP in mTOR cKO microdissected corpus collosum compared to control. Control and mTOR cKO samples were run on the same western blot. Expression of proteins was normalized first to β-tubulin, then to control. Graph presents data from 4 control and 4 mTOR cKO animals. **b**, MOG expression is dysregulated in mTOR cKO corpus callosum. Representative images of MOG staining shows decreased area of MOG positivity in mTOR cKO corpus callosum compared to control. Scale bar = 500 μm. Graph presents data from 8 control and 6 mTOR cKO animals. All values expressed as mean ± SEM *p < .05; **p < .01; ***p ≤ .001. **c**, Demyelination apparent by 12 weeks in the corpus callosum of mTOR cKO mice. Representative EM images from 12-week control and mTOR cKO corpus callosum, scale bar = 1 μm. **d**, Quantification of the number of myelinated axons in 12-week control and mTOR cKO callosa, n = 3/group. **e**, G-ratio scatter plot showing linear regressions for 12-week control (solid) and mTOR cKO (dashed) g-ratios, slope p = 0.048, n > 600/group. **f**, G-ratio distribution of 12-week control and mTOR cKO g-ratios, n > 600/group. **p ≤ 0.01, ****p ≤ 0.0001.

The demyelination that occurs in the brain of the mTOR cKO mice from 8-12 weeks could be due either to adult myelin maintenance deficits or to developmental alterations that do not manifest until after 8 weeks. To determine an effect on myelin maintenance, we deleted mTOR specifically from mature oligodendrocytes using inducible *Plp-Cre^ERT^;mTOR^fl/fl^* adult mice. Myelin analyses revealed deficient long-term myelin maintenance 12 months after mTOR deletion but did not entirely recapitulate the more dramatic phenotype observed in Cnp-mTOR cKO brains at 12 months of age (Supplementary Fig. 9). These data indicate that mTOR likely has distinct and necessary functions in brain oligodendroglia during development and in adult myelin maintenance.

### Developmental loss of mTOR disrupts nodal structures and axonal function in the adult corpus callosum

Demyelination in diseases such as multiple sclerosis (MS) results in progressive neuronal degeneration and functional deficits. Given the significant myelin loss detected in 12-week corpus callosum, we hypothesized that mTOR cKO mice would have compromised axonal function. To determine whether mTOR cKO brains have disrupted nodal structures, we performed immunostaining for contactin-associated protein 1 (CASPR), a component of paranodes, and NaV1.6, a voltage-gated sodium channel present in nodes of Ranvier (Fig. 7a). We observed a significant reduction in the number of paranode-node structures (defined as CASPR-NaV1.6-CASPR) in the midline callosum of mTOR cKO compared to control, suggesting that axonal function may be altered with mTOR loss in oligodendrocytes. To address this directly, we performed compound action potential (CAP) recordings in the midline callosum of coronal slices from 13-week mTOR cKO and control brains (Supplementary Fig. 10). Representative traces revealed clear differences in CAP response (Fig. 7b), with a complete loss of the N1 (myelinated fiber) response and altered N2 (unmyelinated fiber) response in the mTOR cKO callosum. The N1 amplitude was completely absent in mTOR cKO even at the highest stimulation intensity used, while there was a clear N1 response in the control brains (Fig. 7c). While latency to N2 response was unchanged in mTOR cKO corpus callosum (Fig. 7e), we observed an increased duration of the N2 response (Fig. 7f), suggesting changes in the unmyelinated axon response to stimulation. However, the N2 amplitude of the CAP recordings was unchanged across increasing stimulation intensities, indicating that the number of axons is not reduced (Fig. 7d). These data indicate that the endogenous demyelination observed in the mTOR cKO callosum results in significantly impaired myelinated and unmyelinated axonal function in adults, despite the lack of obvious axonal damage (Supplementary Fig. 11).

**Fig. 7.**
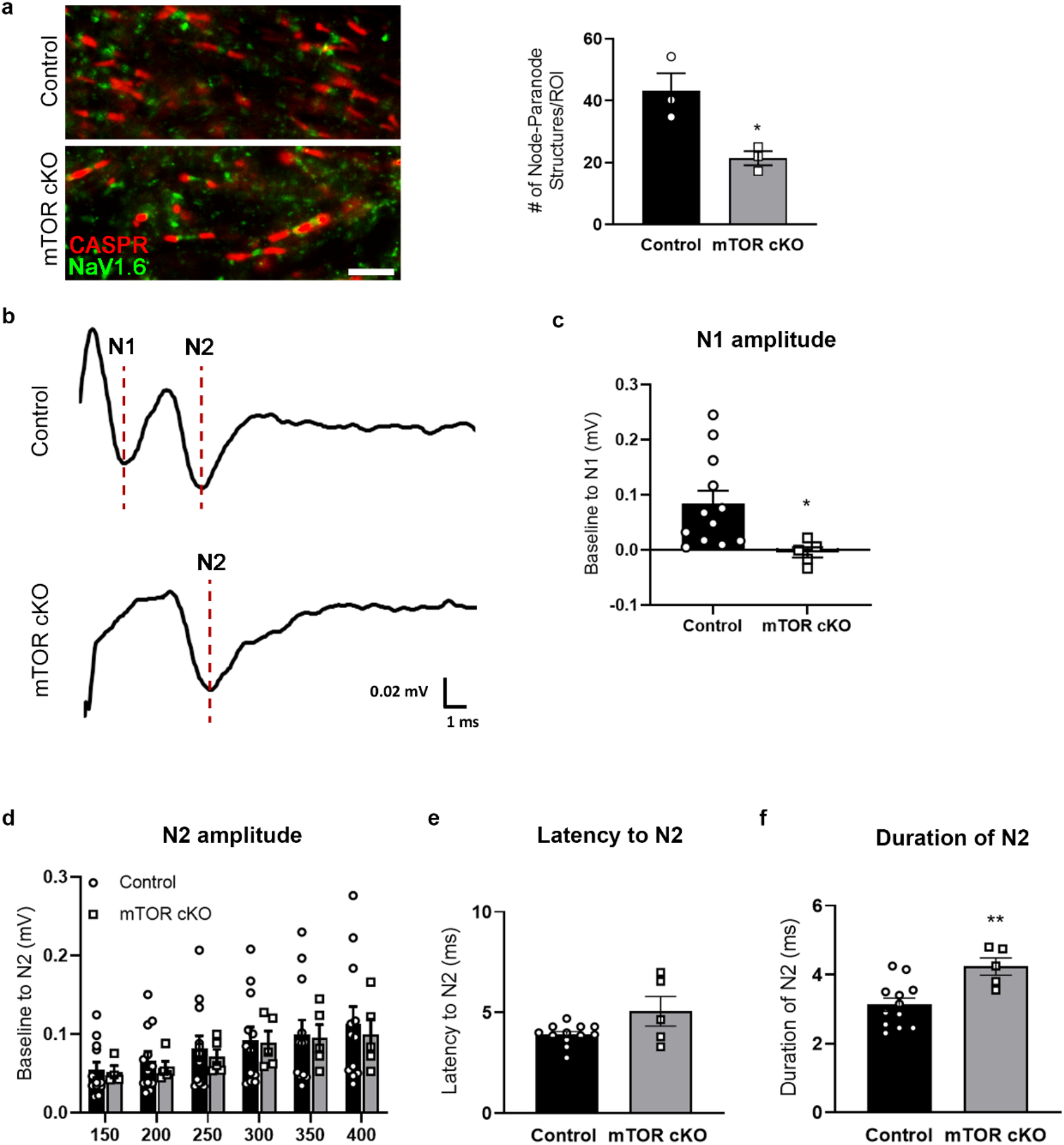
Developmental loss of mTOR disrupts nodal structures and axonal function in the adult corpus callosum. **a**, Representative images of CASPR (red) and NaV1.6 (green) immunostaining in the 12 week control and mTOR cKO callosum, scale bar = 5 μm. Quantification of the number of node-paranode structures (defined as CASPR – NaV1.6 – CASPR) present in the midline callosum, n = 3/genotype. *p ≤ 0.05. **b-f,**Axonal function in the corpus callosum of control and mTOR cKO at 13 weeks of age measured by compound action potential (CAP) callosal slice recordings. **b**, Representative traces from CAP callosal slice recordings. N1, myelinated fiber response. N2, unmyelinated fiber response. **c**, The amplitude of the N1 response at 400μA stimulation intensity, n = 5-12/group. **d**, The amplitude of the N2 response measured across increasing stimulation intensities, n =4-12/group. **e**, The latency to the N2 response at 400μA stimulation intensity, n = 5-12/group. **f**, The duration of the N2 at 400μA stimulation intensity, n = 5-12/group. *p ≤ 0.05, **p ≤ 0.01.

## Discussion

Here we reveal that oligodendrocyte precursors in the brain and spinal cord are transcriptionally distinct. Brain precursors have lower cholesterol biosynthesis compared to spinal cord precursors. mTOR, a major regulator of CNS myelination, promotes cholesterol biosynthesis in oligodendroglia, and loss of mTOR results in differential expansion in brain and spinal cord of a distinct population of oligodendrocytes with reduced cholesterol biosynthesis and dysregulated cytoskeletal dynamics. This demonstrates that mTOR loss has differential effects on distinct oligodendroglial populations. Normal cholesterol biosynthesis is necessary for oligodendrocyte viability and myelin gene expression, and at 8 weeks of age, along with continued reduction of cholesterol biosynthesis, there is increased apoptosis resulting in fewer brain oligodendroglia in mTOR cKO mice. By 12 weeks of age mTOR cKO mice exhibit callosal demyelination and impaired axonal function.

Cholesterol is essential for normal myelination in the CNS^32,33^; so why do brain oligodendroglia have lower cholesterol biosynthesis than spinal cord oligodendroglia? Cells have two sources of cholesterol: *de novo* synthesis and uptake from the extracellular environment. Brain oligodendroglia may have a greater capacity for cholesterol uptake than spinal cord oligodendroglia. Evidence for this comes from our sequencing data showing that *Low-density lipoprotein receptor-related protein-1 (Lrp1),* which functions in receptor-mediated uptake in oligodendroglia^34^, has higher expression in brain precursors than in spinal cord precursors (logFC = 0.82; additional supplementary files), indicating potential for greater cholesterol uptake. Another possible scenario is that brain oligodendroglia require less cholesterol than spinal cord oligodendroglia. Axons are thicker in spinal cord white matter than in brain corpus callosum, requiring proportionally thicker myelin with longer internodes^35–37^, and there is evidence that longer sheath length is a cell-intrinsic property of spinal cord oligodendrocytes^1^. It is noteworthy that the premyelinating oligodendroglia have transcriptomic differences between brain and spinal cord, suggesting that these differences precede effects due to eventual differences in number or length of internodes in the two regions after myelination is complete.

Can the different developmental phenotypes in the mTOR cKO brain and spinal cord be explained by our current findings? At the single-cell level in mTOR cKO mice, we discovered a population of myelin-forming oligodendrocytes that have deficits in cholesterol biosynthesis and cytoskeletal dynamics. This population has a greater expansion in the mTOR cKO spinal cord than the brain, and therefore may contribute to the more pronounced developmental phenotype observed in spinal cord of mTOR cKO^8^. We have shown that spinal cord oligodendroglia normally express higher levels of cholesterol biosynthesis enzymes than brain oligodendroglia, and when mTOR is lost, there is a greater cholesterol deficit in the spinal cord than in the brain. Whether brain oligodendroglia have a greater capacity for cholesterol uptake, or simply a lower demand for it, differential cholesterol synthesis may explain why mTOR loss has less impact on brain myelination than spinal cord myelination.

We considered the prospect of increased cholesterol availability rescuing the mTOR cKO myelin phenotype. Since cholesterol does not cross the blood-brain-barrier^38^, oligodendroglia cannot uptake dietary cholesterol and availability is limited to cholesterol produced by neighboring cells. In order to deliver cholesterol to oligodendroglia through diet, use of an animal model with blood-brain-barrier disruption is necessary^33,39^; therefore we were unable to attempt myelin rescue through a high cholesterol diet.

How does mTOR promote cholesterol biosynthesis in oligodendroglia? In Schwann cells as well as in other mammalian cells types, mTORC1 activates sterol regulatory element-binding proteins (SREBPs) which are master transcriptional regulators of cholesterol biosynthesis^40,41^. mTOR regulation of cholesterol biosynthesis in oligodendroglia may occur through similar mechanisms. While our data place mTOR upstream of cholesterol biosynthesis, studies in zebrafish additionally place mTOR downstream of cholesterol, mediating cholesterol-dependent myelin gene expression^42^.

Increased cholesterol availability has been shown to promote remyelination in mouse models of demyelinating diseases^43^. Our data suggest that cholesterol biosynthesis through mTOR signaling has therapeutic implications in demyelinating diseases. Our findings also reveal the importance of elucidating the role of cholesterol biosynthesis in adult myelin maintenance as this may have implications for long-term use of cholesterol-lowering drugs in humans. Indeed, there is evidence that patients taking statins, which can cross the blood-brain barrier, have lower fractional anisotropy, indicating deficits in white matter integrity, than statin-untreated individuals^44^. Further studies will be necessary to determine neurological effects of long-term use statin treatment.

## Methods

### Mice

All animal protocols were conducted in accordance with the guidelines set forth by Rutgers University and are in compliance with Institutional Animal Care and Use Committee (IACUC) guidelines and the National Institutes of Health (NIH) guidelines for the care and use of laboratory animals. All mice were bred and housed in a barrier facility with a 12-hour light/dark cycle. Mice carrying a *floxed-mTOR* allele^45–47^*(Mtor^tm1.1Clyn^*) were provided by Dr. Christopher Lynch (Pennsylvania State College of Medicine, Hershey, PA). The *floxed-mTOR* mice were bred with a dual-reporter mouse line *(B6.129(Cg)-Gt(ROSA)26Sortm4^(ACTB-tdTomato,-EGFP)Luo^/J,* The Jackson Laboratory). These mice contain the *mT/mG* reporter, which globally expresses tomato red, and, in cells with active Cre-recombinase, express GFP. *CNP-Cre* mice^48^ were obtained from K. Nave. All strains were on a C57/Bl/6 background. Mice homozygous for *mTOR floxed* and *mT/mG* reporter alleles and heterozygous for *CNP-Cre* were used for breeding to generate Cre+ or Cre-littermates for experiments. The resultant *mTOR^fl/fl^* mice expressing Cre *(Cre+/-)* have been previously described^8^ and are hereafter referred to as mTOR cKO with *mTOR^fl/fl^/Cre-/-*littermates used as controls. Both males and females were used in all analyses. For all experiments P0 designated the day of birth.

### Isolation of O4+ cells and efficiency analysis

Control (Cre-/-) and mTOR cKO littermates were used at P14 for O4+ cell isolation from brains and at P10 for isolation from spinal cord (Supplementary Fig. 2a). Mice were rapidly decapitated and spinal cords or brains were dissected. Brain dissection at P14 included forebrain and midbrain, and excluded olfactory bulb, cerebellum, and brainstem. Spinal cord dissection at P10 included whole spinal cord. Tissues were dissociated into single cells using Neural Dissociation Kit (P) (Miltenyi). Dissociated cells from control mice were pooled together and mTOR cKO mice were pooled together. O4+ oligodendroglia were isolated by magnetic-activated cell sorting using anti-O4 microbeads (Miltenyi) and eluted into PBS with 0.5% BSA. Isolation efficiency was analyzed by flow cytometry of the negative and positive isolation fractions (Supplementary Fig. 2b). Cells were incubated with FcR blocking reagent and then stained with O4-APC antibody (1:10, Miltenyi) by incubating for 10 min at 4°C. Cells were then washed twice in PBS with 0.5% BSA and 2 mM EDTA and run on LSR II flow cytometer (BD Biosciences). Data were analyzed using FlowJo software. Isolation efficiency was determined to be ~90% by flow cytometry (Supplementary Fig. 2b), and mRNA expression of oligodendrocyte-specific genes was consistent with previously reported characterization of O4+ cells^49^.

### Single-cell sequencing

Droplet-based single-cell partitioning and single-cell RNA-Seq libraries were generated using the 10X Chromium Controller and the Chromium Single-Cell 3’ Reagent v3 Kit (10X Genomics, Pleasanton, CA) as per the manufacturer’s protocol. Briefly, live O4+ cells in single cell suspension at a concentration of 600 cells/μL were mixed with RT reagents and loaded onto a Single-Cell 3’ Chip along with Gel beads and Partitioning oil in the recommended order and then the chip was processed through 10X Chromium Controller for the generation of Gel Beads-in-Emulsion (GEMs). GEM generation was followed by 3’ Gene Expression Library prep protocol that includes Reverse Transcription, cleanup, cDNA amplification, fragmentation, end repair & A-tail prep, adapter ligation and incorporation of i7sample indices into finished libraries, which are compatible with Illumina next-generation sequencing platforms. Sample quantification and quality control were determined using Qubit Fluorometer (Invitrogen, Life Technologies) and TapeStation (Agilent Technologies, Santa Clara CA) respectively. cDNA libraries were sequenced on Illumina NextSeq500 sequencer (Illumina, San Diego, CA) with a configuration of 26/8/0/98 cycles (Read1-10XBarcode+UMI/i7index/i5index/ Read2-mRNA reads). Single-cell sequencing was done at the Genomics Center, Rutgers New Jersey Medical School and sequencing on the NextSeq 500 was supported by National Institutes of Health grant (S10 OD018206).

### Sequencing data pre-processing and quality control

The cellranger mkfastq pipeline (Cell Ranger Version 3.0.2) was used to process the sequencing run generated as described above, and single-cell gene expression was quantified using cellranger count pipeline. Subsequent QC and analysis were performed using Seurat package version 3.0.2 within R 3.6.0. Cells with mitochondrial gene counts >5% were removed. To minimize potential ambient RNA or duplets/multiplets, cells with transcript numbers between 500 and 3000 were selected for downstream analysis. Contaminating non-oligodendroglial cells were eliminated based on known gene markers. After filtering, a total of 9325 cells from spinal cord (3879 control cells, 5446 mTOR cKO cells) were obtained, and a total of 16397 cells from brain (8220 control cells, 8177 mTOR cKO cells) were obtained.

### Data normalization, clustering, and integration

Data were normalized using the default scaling factor of 10000. Principal component analysis was performed prior to clustering and the first 17 principal components (PCs) were used based on the ElbowPlot, with top 2,000 most variable genes. Clustering was performed using the FindClusters function with a resolution of 0.5. The non-linear dimensional reduction and UMAP analysis were performed using the same PCs as input to the clustering analysis. With 2 dimensional reductions, both PC analysis and UMAP calculated, the 4 samples were integrated together using a computational strategy previously described^50^. A few remaining non-oligodendroglial cell clusters were identified based on expression markers and were removed, resulting in the final dataset of 4603 brain control cells, 3887 brain mTOR cells, 3605 spinal control cells, 5233 spinal mTOR cells, which were re-integrated and re-clustered for the downstream analysis.

Similarly, data from O4+ cells from control brain and spinal cord were integrated together and UMAP analysis was performed with the top 20 PCs. A total of 4509 cells from control brain, and 3571 cells from control spinal cord were included for reintegration and re-clustering for final analysis.

### Analysis of differential gene expression

Differential expression of genes between conditions or clusters was performed using the FindMarker function of the Seurat package, which identifies negative and positive markers for each condition or cluster. A gene is required to be detected at a minimum 20-25% in either of the two groups of cells. All genes demonstrating statistically significant differential expression were included in downstream IPA analysis.

### Analysis of frequency and proportion equality

The number of cells in each cluster was examined and percentage of each sample in different clusters was analyzed using two-proportions z-test for equality of proportions. A classical P value was calculated to evaluate the significance.

### Analysis of trajectory and pseudotime

The monocle3 version 0.2.1 within R 3.6.3 was used for trajectory and pseudotime analysis. Cells from the 4 samples integrated in Seurat were used to construct the monocle object, and the Seurat clustering information was preserved to facilitate the interpretation of the pseudotime analysis. The method of UMAP was used to reduce the dimensionality of this dataset, followed by clustering, that starts the construction of trajectory. Cells were ordered according to its progress along a learned trajectory, which is termed as “pseudotime” in monocle. Clusters 8 and 12 were manually identified as OPCs (Supplementary Fig. 3) using the top 5 differentially expressed genes and matched to two reference databases^5,14^. These are the least differentiated clusters and were therefore selected as the root group for pseudotime analysis.

### Functional analysis of differential gene expression

Differential gene expression data were further analyzed using IPA (QIAGEN, https://www.qiagenbioinformatics.com/products/ingenuity-pathway-analysis). Network analysis in Supplementary Fig. 4a was generated using IPA.

### Protein extraction and immunoblotting

For analyses at P10, P14, and P18, O4 cells were isolated and lysed in RIPA lysis buffer (Thermo Scientific, 89900) with 1x Halt Protease/Phosphatase Inhibitor Cocktail (Thermo Scientific, 78440). For analyses at 8 weeks and 12 weeks, brains were dissected from control and mTOR cKO mice. Brains were cut into 2 mm wide coronal slices directly above the hippocampus using a brain mold. The midline corpus callosum was microdissected from the coronal slice and immediately frozen in Eppendorf tubes on dry ice. To prepare protein samples, frozen samples were ground into a fine powder using a micro-mortar and pestle. Samples were then lysed using 60 μL RIPA buffer with 1x Halt Protease/Phosphatase Inhibitor Cocktail. Samples were gently pipetted for 1 min to homogenize.

The RC-DC protein assay (BioRad) was performed to determine protein concentration. 20 μg of protein was resolved by SDS-PAGE on 4-12% Bis-Tris mini-gels (Invitrogen). Separated proteins were transferred to nitrocellulose membranes and blocked in 5% milk/TBS-0.1% Tween for 2 hr at room temperature. Membranes were then incubated in the presence of primary antibodies diluted in 5% BSA/TBS-0.1% Tween overnight at 4°C. The exception was anti-FDPS which was incubated in 5% milk/TBS-0.1% Tween for 1.5 hr at room temperature. Membranes were then washed 3 times for 10 min with TBS-0.1% Tween and incubated for 1 hr at room temperature in 5% milk/TBS-0.1% Tween containing goat anti-rabbit or goat antimouse HRP-conjugated secondary antibodies at a dilution of 1:5000. The detection of HRP conjugated secondary antibodies was performed by enhanced chemiluminescence using the Ultra-LUM imaging device. Protein expression levels were quantified using Image J. The following antibodies were used: Hmgcr (Invitrogen, PA5-37367 1:500), Hmgcs1 (Cell Signaling, 42201S, 1:1000), Fdps (Proteintech, 16129-1-AP, 1:1000), β-actin (Sigma, A5441, 1:5000), MBP (Covance, SMI-99, 1:500), PLP (Abcam, ab28486, 1:3000), β-tubulin (Cell Signaling, 5346S, 1:1000), and β-amyloid (Invitrogen, 51-2700, 1:1000), SMI-32 (Biolegend, 801701, 1:500). Statistical analysis was performed using GraphPad Prism. Unpaired two-tailed t test was performed to determine significance.

### Primary rat OPC cultures

OPCs were purified from cortical mixed glial cultures by established methods^51^. Briefly, brains were removed from postnatal day 0–2 Sprague Dawley rat pups and the cortices were dissected. Cortical pieces were enzymatically digested in 2.5% trypsin and Dnase I followed by mechanical dissociation. Cells were resuspended in MEM-C, which consisted of minimal essential media (MEM) supplemented with 10% FBS, L-glutamine, and 1% Pen-strep, and plated in T75 flasks. The resulting mixed glial cultures were maintained until confluent for a day (~10 days total). Purified OPC cultures were prepared by a differential shake. Purified OPCs were seeded onto poly-D-lysine coated T75 flasks at a density of 1.5 × 10^4^ cells/cm2 in N2S media. N2S consists of 66% N2B2 (DMEM/F12 supplemented with 0.66 mg/ml BSA, 10 ng/ml d-biotin, 5 μg/ml insulin, 20 nM progesterone, 100 μM putrescine, 5 ng/ml selenium, 50 μg/ml apotransferrin, 100 U/ml penicillin, 100 μg/ml streptomycin, and 0.5% FBS) supplemented with 34% B104 conditioned media, 5 ng/ml FGF, and 0.5% FBS. To initiate oligodendrocyte differentiation, we followed an established mitogen withdrawal protocol^52^. After overnight recovery in N2S, media was replaced with Differentiation media (mitogen free N2B2 media supplemented with 30 ng/ml triiodothyronine (T3)). Differentiation media with or without inhibitors was replenished every 48 hr over the course of experiments. Cell culture media (MEM, DMEM/F12), FBS, trypsin, and insulin-selenium-transferrin (ITS) were purchased from Gibco (Long Island, NY). Additional N2 supplements, T3, and poly-d-lysine were purchased from Sigma. Recombinant human FGF-2 was purchased from R&D Systems (Minneapolis, MN).

### Treatment with inhibitors

To inhibit mTOR, cells were treated with 10 nM rapamycin (Calbiochem, 553210) which is an effective dose based on our prior studies^22,53^. Stock solutions of 5mM rapamycin were prepared in DMSO. Control cultures received vehicle alone.

To inhibit cholesterol biosynthesis, cells were treated with simvastatin (Millipore, 567022). Stock solutions of 1mM simvastatin were prepared in DMSO. While the effective dose of simvastatin for HMGCR inhibition in oligodendroglia *in vitro* is unclear, the IC80 for cholesterol synthesis in rat hepatocytes is approximately 5 μM simvastatin^54^. Therefore, we treated differentiating primary rat OPCs with 5 μM and 1 μM simvastatin. Control cultures received vehicle alone.

### RNA isolation and reverse transcription

RNA was extracted using RNeasy Plus Mini Kit (Qiagen, Valencia, CA). RNA concentration was measured using a NanoDrop spectrophotometer (Thermo Scientific) and cDNA was reverse transcribed using Superscript II (Invitrogen).

### Quantitative real-time PCR

RT-PCR was performed using SYBR green detection master mix (BioRad 430001607) and amplification was normalized to expression levels of Gapdh for each sample. β-actin was used as an additional housekeeping control to confirm that Gapdh expression was constant in treated oligodendroglia in all conditions. RT-PCR was performed on the Applied Biosystems 7900B (Carlsbad, CA) using the associated Sequence Detection System Version 2.2.2. The thermal reaction profiles were performed as previously described^53^. BioRad plates and QuantiTect primers were used. Primers (Qiagen): Gapdh QT00199633, Mbp QT00199255, Mag QT00195391, Plp1 QT00176414, Hmgcr QT00182861, Hmgcs1 QT00183267, Fdps QT00175574, β-actin QT00193473. Statistical analysis for each target was performed using unpaired twotailed t test (Fig. 4c) or one-way ANOVA followed by Tukey’s post hoc multiple comparisons test (Fig 5) in GraphPad Prism.

### Cholesterol analysis

Brain and spinal cord tissue were dissected from mice at P14 and P10, respectively and O4+ cells were isolated as described above. Neutral lipids were extracted using a modified Folch method^55^. Briefly, cells were homogenized in chloroform:methanol (2:1) and incubated at room temperature for 1 hour. Following addition of 0.9% sodium chloride, each sample was centrifuged at 2000xg, 4°C for 10 minutes. The organic phase was removed, dehydrated overnight, and resuspended in a butanol:Triton-X:methanol solution. Cholesterol content was determined using an enzymatic colorimetric cholesterol assay (StanBio Laboratory, Boerne, TX) and normalized to cell count. Unpaired two-tailed t test was performed to determine significance using GraphPad Prism.

### Cell viability assay

Cell viability was measured after 3 days and 5 days of culture using an MTT [3-(4,5-dimethyldiazol-2-yl)-2,5-diphenyl tetrazolium bromide] assay (Abcam, ab211091) as per manufacturer’s protocol. Briefly, cells were incubated with MTT reagent for 3 hours at 37°C before adding MTT solvent and mixing on an orbital shaker for 15 mins. Absorbance was measured at OD590 nm and results were normalized to control culture conditions. Each condition was plated and analyzed in triplicate and the experiment was repeated three times. Statistical analysis for was performed using one-way ANOVA followed by Tukey’s post hoc multiple comparisons test in GraphPad Prism.

### Tissue preparation and immunostaining

mTOR cKO animals and corresponding controls were taken for analysis at 7-8 weeks and 12 weeks of age. A total of four to six animals per genotype from the corresponding time point were used for all analyses described. Mice were intracardially perfused with 10 mL of an icecold PBS solution containing phosphatase inhibitors and heparin followed by 40 mL of ice-cold 3% paraformaldehyde (PFA) at a rate of 2 mL/min. Brains were dissected and cut into 2 mm wide coronal slices directly above the hippocampus using a brain mold and drop-fixed in 3% PFA overnight. The tissue was then dehydrated with a 30% sucrose solution and subsequently frozen in OCT. 20 μm serial sections were taken throughout the brain slices. Immunolabeling was performed as described previously^56^, with specific treatments used for antibody staining as necessary. Briefly, for myelin oligodendrocyte glycoprotein (MOG) staining, sections were delipidated for 10 min in 100% ethanol then treated with 3% hydrogen peroxide to eliminate endogenous peroxidase activity. The NovaRed substrate kit (SK-4800, Vector Laboratories) was used to detect positive signal, then slides were dehydrated in ethanol and coverslipped with Cytoseal (8312-4, Thermo Scientific). For SOX10 and cleaved caspase-3 staining, sections were soaked for 10 min in 0.01 M sodium citrate buffer heated to boiling before immersion. Sections were blocked for 60 min at room temperature in 3% donkey or goat serum (D9663, Sigma Aldrich) in 1x PBS. Sections were incubated at 4°C overnight in primary antibodies diluted in 1% donkey serum, and 0.4% Triton X-100 (T8787, Sigma Aldrich) in 1x PBS. Primary antibodies used: goat anti-SOX10 (1:50, AF2864, R&D Systems), rabbit anti-cleaved caspase-3 (1:400, 9664S, Cell Signaling), rabbit anti-MOG (1:1000, ab32760, Abcam), mouse anti-CASPR (1:200, MABN69, Millipore), and rabbit anti-NaV1.6 (1:300, ASC-009, Alomone Labs). Secondary antibodies used: donkey anti-mouse 647 (1:500, A31571, Life Technologies), donkey anti-goat AMCA (1:100, 705-155-147, Jackson Laboratories), donkey anti-rabbit 647 (1:750, A31573, Life Technologies), goat anti-mouse 647 (1:500, A21235, Life Technologies), and biotinylated goat anti-rabbit (1:500, BA-1000, Vector Laboratories) followed by streptavidin-HRP (1:1000, 21126, Pierce). Immunofluorescent stained sections were coverslipped using Fluorogel (17985-10, Electron Microscopy Sciences) then sealed with nail lacquer.

### MOG image analysis

To analyze MOG immunostaining, images were taken with a 2x objective using an Olympus AX-70 microscope and analyses were performed using ImageJ software. The entire midline callosum visible in each 2x image was outlined as the ROI. MOG positivity was defined using the adjust threshold function for positive staining and the area fraction within the ROI was calculated using ImageJ software. Data were then expressed as a percentage of the total ROI. At least 3-4 images/animal and 3-8 animals/group were analyzed. Unpaired two-tailed t test was performed to determine significance using GraphPad Prism.

### Cell counts

To analyze SOX10/cleaved caspase-3 immunostaining, images were taken with a 20x objective using an Olympus AX-70 microscope. Cell counts were performed using ImageJ software with the cell counter plugin. The midline callosum visible within the image was outlined as the ROI. The area in μm^2^ for each ROI was used to normalize all data to mm^2^. For all counts, 3-4 images/animal and 6 animals/group were analyzed. All SOX10+ cells within an ROI were counted, followed by all double-immunolabeled cleaved caspase-3+/SOX10+ cells. To analyze CASPR and NaV1.6 immunostaining, images were taken with a 60x oil objective using an A1R confocal microscope. Counts of nodal structures, defined as two CASPR+ paranodes on either side of a NaV1.6+ node, were performed using ImageJ software and the cell counter plugin. The entire image (2621.44 μm^2^) was used as the ROI. For all counts, 3-4 images/animal and 3 animals/group were analyzed. Unpaired two-tailed t test was performed to determine significance using GraphPad Prism.

### Electron microscopy

Mice were perfused and brains were dissected as described above. A 2 mm region just rostral of the hippocampus was drop-fixed in 2% PFA/2.5% glutaraldehyde (01909-100, Polysciences, Inc.) at 4°C overnight. Using a PELCO Biowave Pro tissue processor (Ted Pella), the tissue was rinsed in 100 mM cacodylate buffer and then post-fixed in a reduced osmium mixture consisting of 1% osmium tetroxide and 1.5% potassium ferrocyanide followed by 1% osmium tetroxide alone. Dehydration was carried out in a graded series of acetone (50%, 70%, 90%, 100%) containing 2% uranyl acetate for en bloc staining. Finally, tissue was infiltrated and embedded in Embed 812 (Electron Microscopy Services) and cured for 48 hr at 60°C in an oven. The corpus callosum pieces were oriented such that sections could be cut midline in a sagittal plane. Ultrathin sections (65 nm) were mounted on copper mesh grids and viewed at 80 kV on a Tecnai G2 transmission electron microscope (FEI). Electron micrographs of the corpus callosum were imaged near the midline.

In order to calculate g-ratios, all axons within 3 electron micrographs/animal were measured using ImageJ software to calculate the area of each axon as well as the total area of the myelinated fiber. Axon diameter and total myelinated fiber diameter were then extracted from each measured area in Microsoft Excel. At least 199 and up to 565 axons were measured in each animal with 3 animals/group. G-ratio was calculated as axon diameter/total myelinated fiber diameter. For scatter plots and g-ratio distribution graphs, all measured g-ratios were plotted. In order to examine number of myelinated axons, all myelinated axons within 3 electron micrographs/animal were counted using ImageJ software and the cell counter plugin. The number of myelinated axons within each animal was defined as the average number of myelinated axons in each micrograph. The ROI was defined as the entire micrograph area, 141.23 μm^2^. For g-ratio scatter plots, linear regression analyses were used to compare the slope and y-intercept of the control and Cnp-mTOR cKO or Plp-mTOR cKO linear regressions using all measured g-ratios. For g-ratio distribution analysis, the Kolmogirov-Smirnov t-test was used in order to compare cumulative ranks using all measured g-ratios.

### Callosal slice electrophysiology

Electrophysiology recordings were performed *in vitro* on coronal slices of control and mTOR cKO brains. Slicing ACSF (125mM NaCl, 2.4mM KCl, 2mM MgCl_2_, 1.015 mM NaH_2_PO_4_.H_2_O, 26.40mM NaHCO3, 2.04mM CaCl_2_.2H_2_O, and 11mM Glucose) and recording ACSF (125mM NaCl, 2.4mM KCl, 1.4mM MgCl_2_, 1.015mM NaH_2_PO_4_, 26.40mM NaHCO_3_, 2.18mM CaCl_2_.2H_2_O, and 24.94mM Glucose) were prepared. Mice were anesthetized using isoflurane and quickly decapitated. Brains were rapidly dissected and submerged in ice-cold slicing ACSF solution for 3 min. The cerebellum and 1/3 of prefrontal/frontal cortex were separated and the rest of the brain was submerged in the same solution for an additional 3 min. After the brains were adequately cooled, coronal slices of 400 μm thickness were prepared using a LEICA VT 1200S vibratome. The coronal slices were incubated in oxygenated half slicing–half recording ACSF solution (to prevent glucose shock) for 2 hours at 31-32°C. Electrophysiology recordings were done on the corpus callosum. The FHC concentric bipolar stimulation electrode, CBBPE75, was inserted on one hemisphere and the micropipette glass recording electrode filled with 3M NaCl was located on the other hemisphere (Supplementary Fig. 10). The Axon Digidata 1550B and Multiclamp 700B of Molecular Devices were used to acquire and amplify the extracellular signal. Field compound action potentials (CAP) recorded through the recording electrode were analyzed for N1 (myelinated fiber) as well as N2 (unmyelinated fiber) amplitude, latency, and duration. For slice recording analyses, Student’s t-test was performed using all measured recordings from control and Cnp-mTOR cKO groups. All data are presented as mean ±SEM.

## Acknowledgements

We thank Alex Lemenze and Robert Donnelly for initial assistance with single-cell transcriptomics, Quan Shang for technical assistance, and Angelina Evangelou and Andrew Thomas for helpful discussions. We also thank Mainul Hoque and Neeraja Syed at the Rutgers NJMS Genomics Center for sequencing performed on Illumina NextSeq 500 (NIH SIG grant 1S10OD018206-01A1). The authors acknowledge the Office of Advanced Research Computing (OARC) at Rutgers, The State University of New Jersey for providing access to the Amarel cluster and associated research computing resources that have contributed to the results reported here. URL: https://oarc.rutgers.edu. This work was supported by the National Institute of Neurological Disorders and Stroke R01/R37 NS082203 to T.L.W. and W.B.M. and F31 NS108521 to M.A.J. and the National Multiple Sclerosis Society RG5371-A-4 to T.L.W.

## Author contributions

L.K and T.L.W conceptualized the project. L.K, M.A.J, M.L.M, and I.M.O performed experiments and analyzed data. Y.J.C analyzed single-cell sequencing data. J.N.B performed electron microscopy. A.K.T performed callosal slice electrophysiology. L.K and M.A.J prepared the figures. L.K and M.A.J wrote the manuscript. L.K, M.A.J, Y.J.C, M.L.M, I.M.O, O.B.G, W.B.M, and T.L.W edited the manuscript.

---

**Supplementary Fig. 1.**
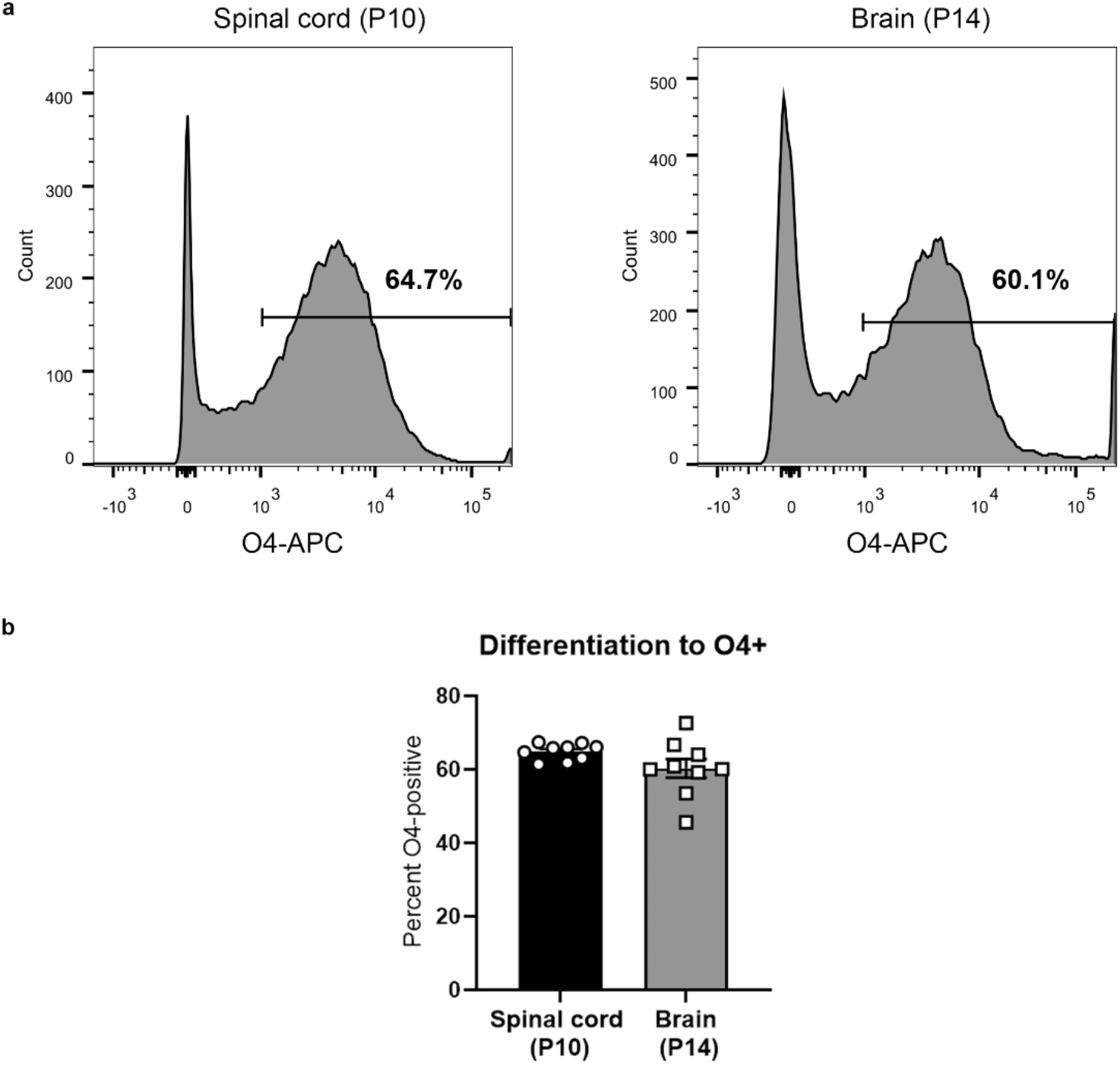
Characterization of differentiation stage in P10 spinal cord and P14 brain. **a**, Spinal cords at P10 and brains at P14 from control mice were dissociated and cell suspensions were analyzed by flow cytometry. O4 was used as a differentiation marker. Representative histograms showing O4+ populations in P10 spinal cord and P14 brain. **b**, Quantification of the percentage of O4+ cells in P10 spinal cord and P14 brain. Values expressed as mean ± SEM in percentage of live cells, n=9/region.

**Supplementary Fig. 2.**
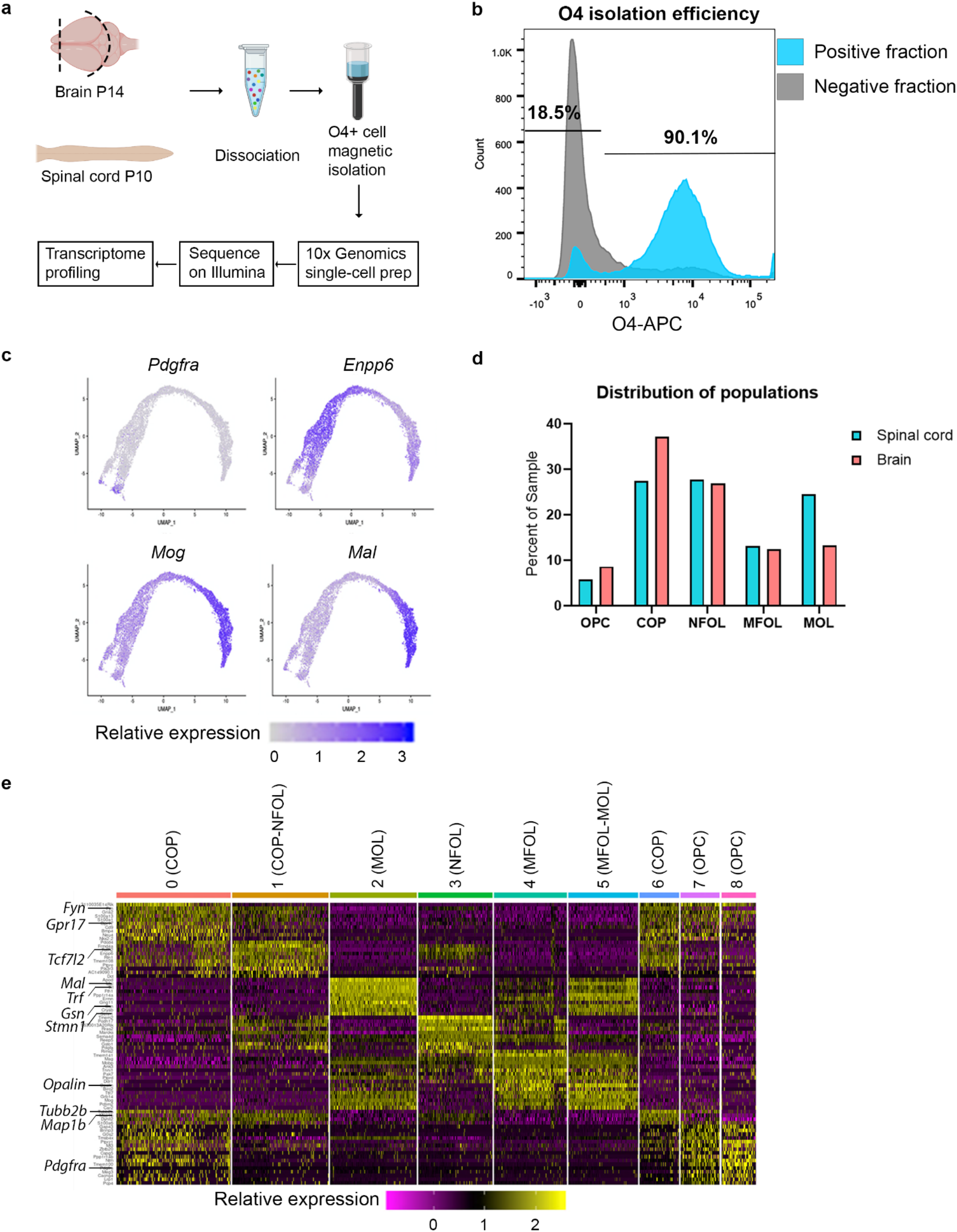
Experimental approach and characterization of O4+ cells. **a**, Brain dissection at P14 included forebrain and midbrain, and excluded olfactory bulb, cerebellum, and brainstem. Spinal cord dissection at P10 included whole spinal cord. Tissues were dissociated and O4+ oligodendroglia magnetically isolated. Single-cell transcriptional profiling was performed using 10x Genomics capture and Illumina sequencing technology. Created with Biorender. **b**, O4+ cell isolation efficiency analyzed by flow cytometry of the negative and positive isolation fractions. **c**, Relative expression of known markers of differentiation. Populations form a differentiation continuum beginning with *Pdgfra+* OPCs, followed by *Enpp6+* cells that are transitioning into newly formed oligodendrocytes, and ending with more mature cell clusters marked by *Mog* in myelin-forming and mature oligodendrocytes, and *Mal* in mature oligodendrocytes. **d**, Single-cell sequencing shows distribution of oligodendroglial populations at each differentiation stage in spinal cord and brain. There is a minor shift towards more precursors (OPCs and COPs) in the brain (12.7% difference), equivalent proportions of NFOLs, and MFOLs in the two CNS regions, and a small shift towards more MOLs in the spinal cord (11.2% difference). **e**, Heatmap demonstrates gene signatures of 9 distinct clusters. Relative expression is shown for top 10 differentially expressed genes in each cluster. Color key shows high and low relative expression. OPC clusters 7 and 8 have high expression of *Pdgfra,* which promotes precursor motility and inhibits differentiation (Noble 1988); COP clusters have downregulated *Pdgfra* and express genes with known functions in early OPC differentiation including *Gpr17* (G protein receptor 17) that regulates timing of oligodendrocyte differentiation (Chen 2009), *Fyn* that functions in cytoskeletal regulation of process outgrowth (Richter-Landsberg 2008) and the microtubule or microtubule-associated genes, *Tubb2b*(beta-tubulin) and *Map1b* (microtubule associated protein 1b). Cells transitioning from COP to NFOL (cluster 1) are characterized by expression of *Tcf7L2* (transcription factor 7 like 2), a transcription factor important for oligodendrocyte differentiation (Weng 2017); MFOL cells have high expression of *Opalin* (oligodendrocytic myelin paranodal and inner loop protein) which promotes process extension and branching (de Faria 2019). MOL cells (cluster 2) express *Mal* and *Trf*, which are important for myelination (Schaeren-Wiemers 2004, Espinosa de los Monteros 1999) and *Gsn* and *Stmn1* that destabilize the actin cytoskeleton or microtubules, respectively (Liu 2003).

**Supplementary Fig. 3.**
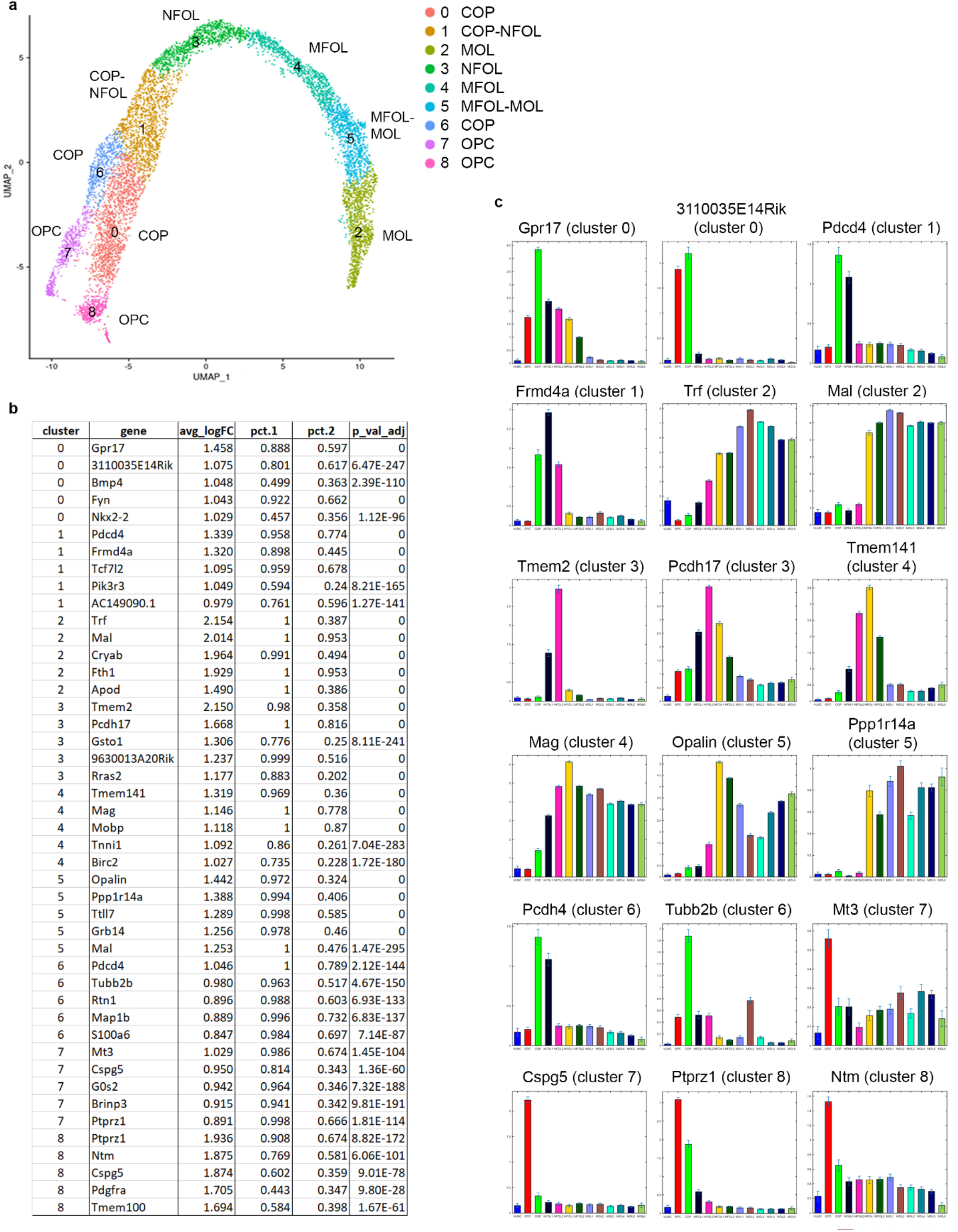
**a**, UMAP of 8,208 O4+ oligodendroglia from developing brain and spinal cord. Cells cluster along a differentiation continuum from oligodendrocyte precursor cells (OPC) to committed OPCs (COP), newly formed oligodendrocytes (NFOL), myelin forming oligodendrocytes (MFOL, and finally mature oligodendrocytes (MOL). 3 littermates pooled for brain; 4 littermates pooled for spinal cord. **b**, Top 5 differentially expressed genes in each cluster compared to all others. avg_logFC=average log fold change; pct.1= percent of cells in cluster expressing listed gene; pct.2= percent of genes in all other clusters expressing listed gene. **c**, Top two genes for each cluster were searched in Castelo-Branco single-cell database (Marques 2016) to identify differentiation stage. X axis labels on all graphs: VLMC, OPC, COP, NFOL1, NFOL2, MFOL1, MFOL2, MOL1, MOL2, MOL3, MOL4, MOL5, MOL6.

**Supplementary Fig. 4.**
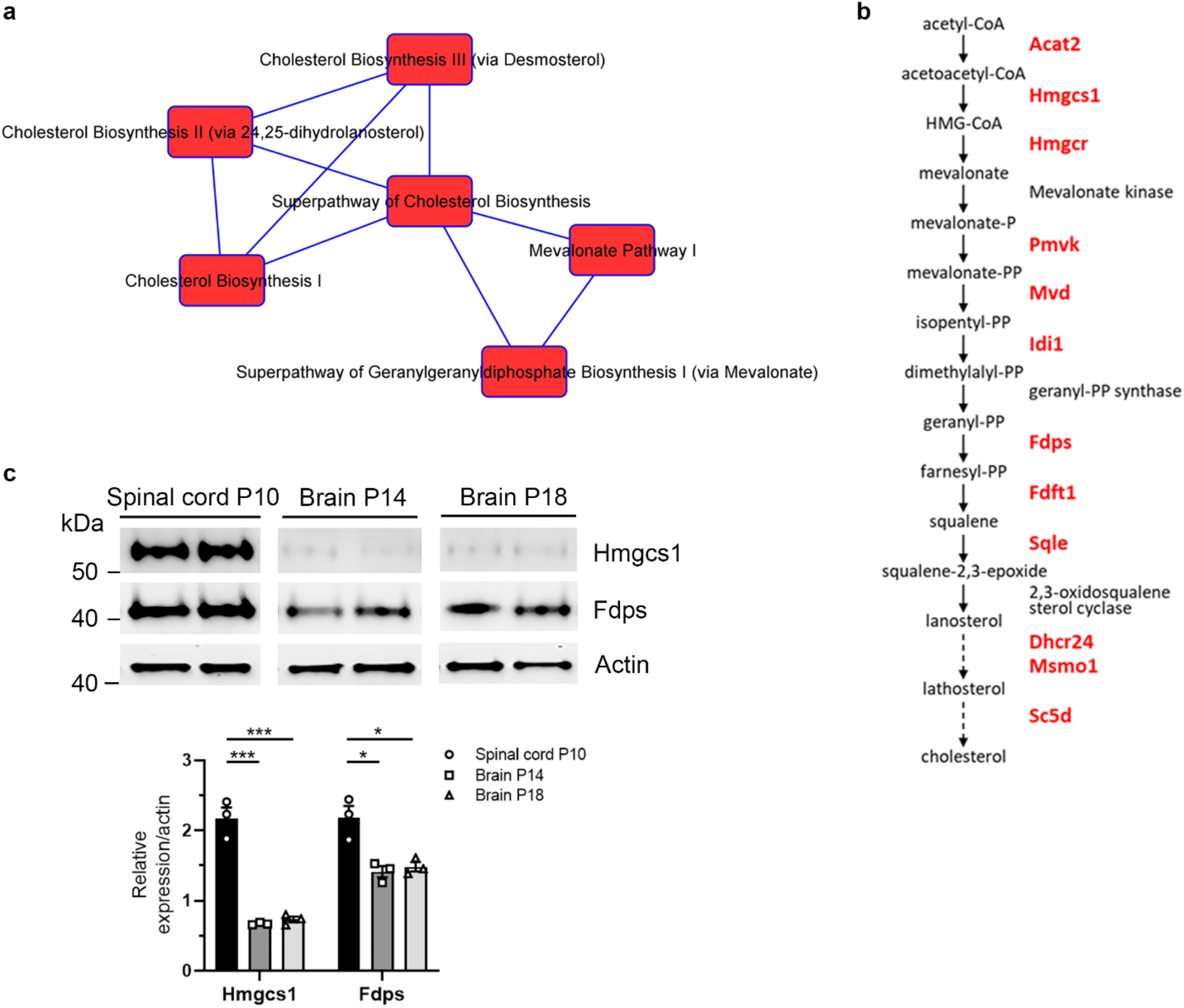
**a**, Top pathways that are differentially expressed in brain (compared to spinal cord) and mTOR cKO (compared to control) are components of, or associated with, cholesterol biosynthesis. Figure was generated in IPA by selecting top canonical pathways to show overlapping pathways. Overlapping genes include Farnesyl diphosphate synthase *(Fdps),* a branch point between downstream cholesterol biosynthesis and geranylgeranyl diphosphate (GGDP) biosynthesis. The mevalonate pathway, cholesterol biosynthesis I, and cholesterol biosynthesis II are all components of superpathway of cholesterol biosynthesis. **b**, Selected substrates, enzymes, and products of cholesterol biosynthesis showing relation to each other within the pathway. Enzymes marked in red have lower expression at the transcript level in brain oligodendroglia compared to spinal cord. **c**, Brain oligodendroglia have lower expression of cholesterol biosynthesis enzymes compared to spinal cord. To rule out the possibility that lower expression of HMGCS1 and FDPS is due to a more immature state of the cells in the brain versus spinal cord, we isolated brain O4+ cells at P18 and found that the brain cells continue to have lower expression of cholesterol biosynthesis enzymes even at a later developmental stage. Representative western blots showing expression of the cholesterol biosynthesis enzymes HMGCR, HMGCS1 and FDPS in O4+ cells isolated from P10 spinal cord, P14 brain, and P18 brain. P10 spinal cord, P14 brain and P18 brain were run on the same western blot. Data quantified from 3 animals/genotype/CNS region. Values expressed as mean ± SEM *p < 0.05; ***p < 0.001

**Supplementary Fig. 5.**
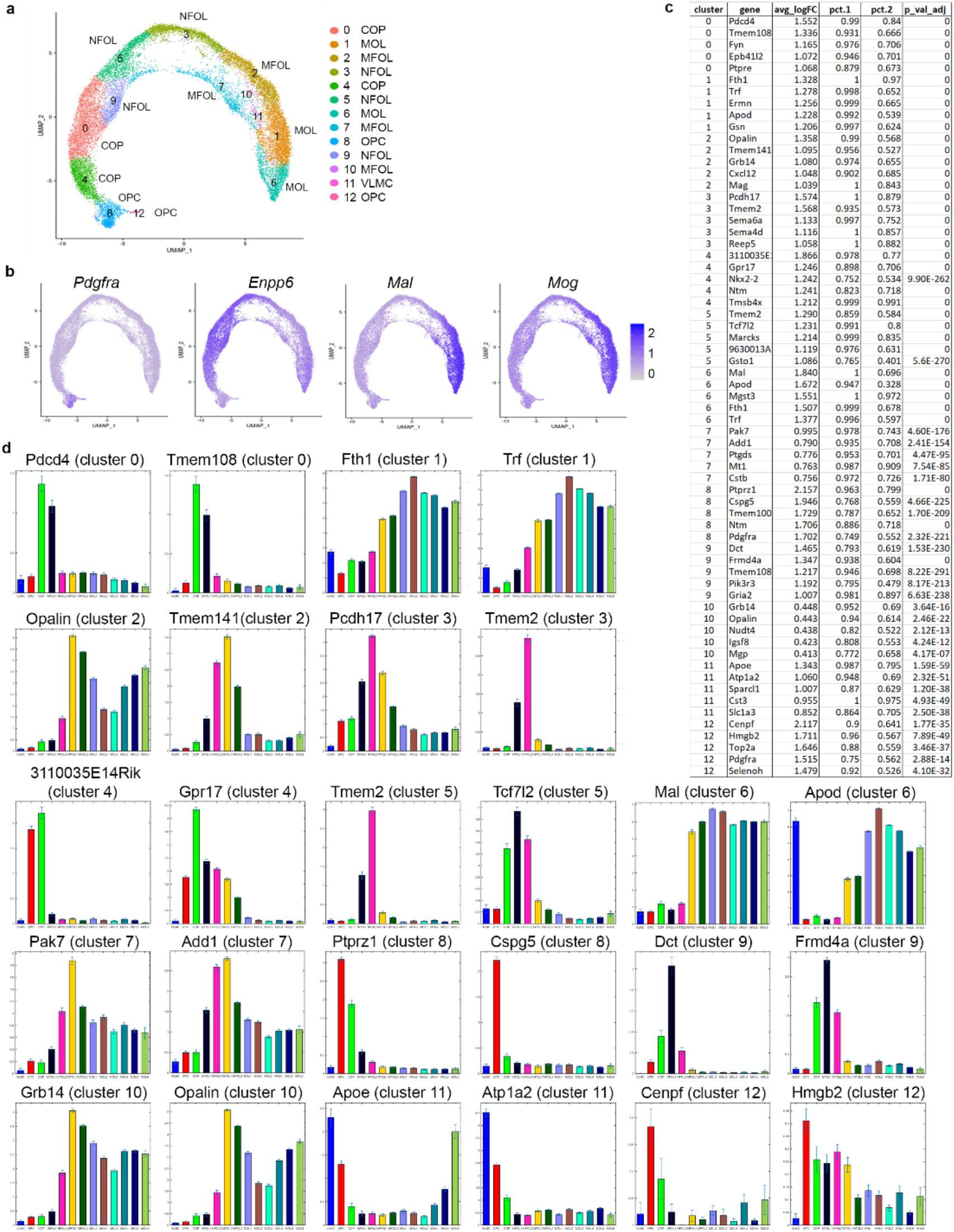
**a**, Combined UMAP of O4+ cells from brain and spinal cord of control and mTOR cKO animals. Cells cluster in a differentiation continuum from OPCs to COPs, NFOLs, MFOLs, and MOLs with one non-oligodendroglial VLMC (vascular leptomenigeal cell) cluster (cluster 11) which was excluded from further analyses. OPCs in cluster 12 have a higher expression of cell cycle genes *(Cdca8, Ccnd2, Cks2, Cdkn3, Cdca3, Cdca7, Cdk6),* indicating that they are cycling cells. 8,208 control cells (3,605 from P10 spinal cord, 4,603 from P14 brain); 9,120 mTOR cKO cells (5,233 from P10 spinal cord, 3,887 from P14 brain). 3 littermates/genotype pooled for brain; 4 littermates/genotype pooled for spinal cord. **b**, Relative expression of known markers of differentiation show that populations form a differentiation continuum beginning with *Pdgfra+* OPCs, followed by *Enpp6+* cells that are transitioning into newly formed oligodendrocytes, and ending with more mature cell clusters marked by *Mog* in myelin-forming and mature oligodendrocytes, and *Mal* in mature oligodendrocytes. **c**, Top 5 differentially expressed genes in each cluster compared to all others. avg logFC= average log fold change; pct.1= percent of cells in cluster expressing listed gene; pct.2= percent of genes in all other clusters expressing listed gene. **d**, Top two genes for each cluster were searched in Castelo-Branco single-cell database (Marques 2016) to identify differentiation stage. X axis labels on all graphs: VLMC, OPC, COP, NFOL1, NFOL2, MFOL1, MFOL2, MOL1, MOL2, MOL3, MOL4, MOL5, MOL6.

**Supplementary Fig. 6.**
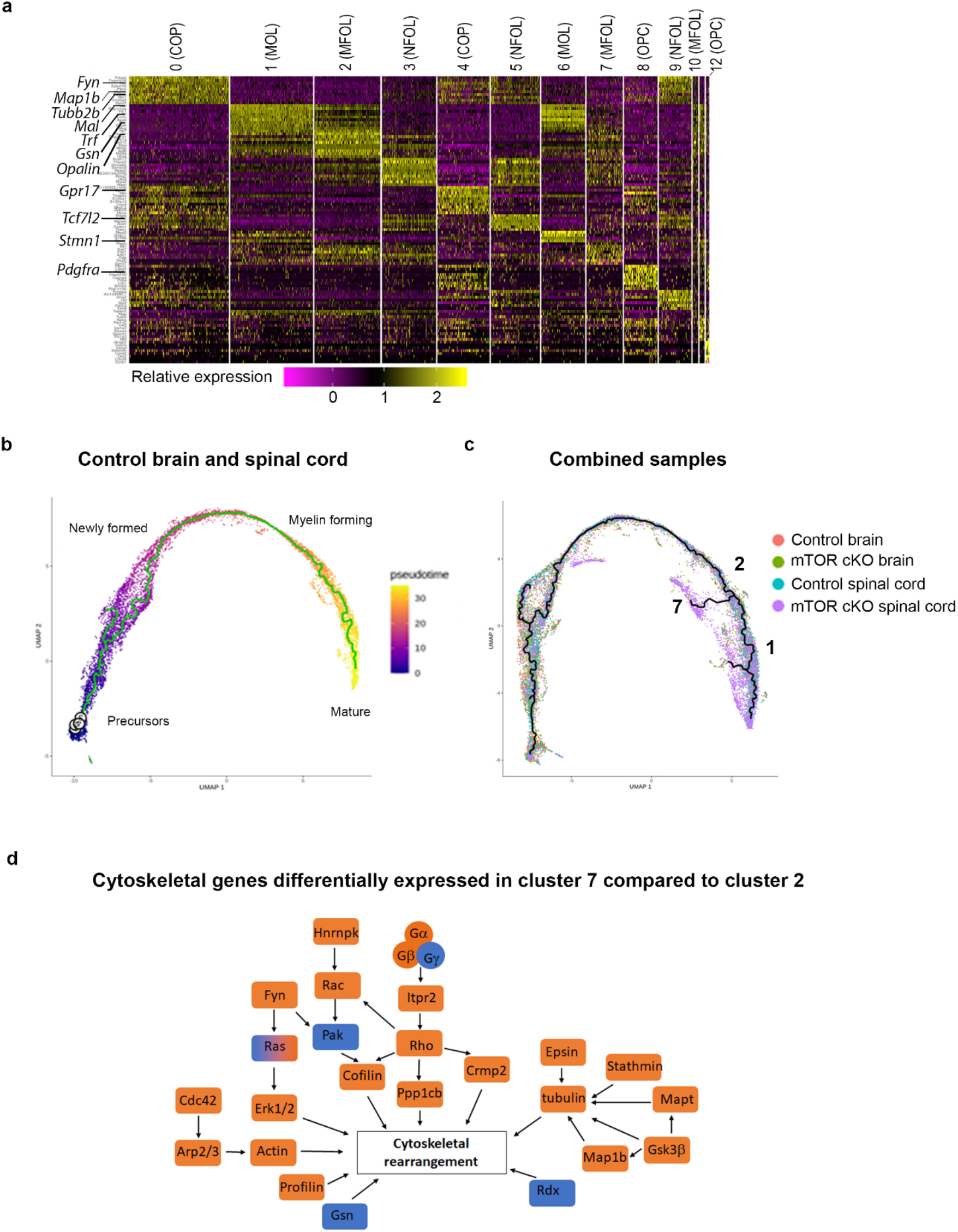
**a**, Heatmap of the top differentially expressed genes in each cluster (combined samples, Fig. 2a, Supplementary Fig. 5a) demonstrates that the regulators of differentiation and myelination noted in Supplementary Fig. 1d are also driving the clustering pattern in the combined UMAP. **b**, Cells from single-cell analysis of control brain and spinal cord are ordered by pseudotime algorithms using Monocle. Trajectory begins at OPCs in purple and progresses to MOLs in yellow. Trajectory follows unbranching pattern between myelin forming and mature oligodendrocytes. **c**, Pseudotime trajectory of combined samples. MFOL cluster 2 branches into MFOL cluster 7, which consists mostly of mTOR cKO cells, and MOL cluster 1 which consists of both control and mTOR cKO cells. **d**, Cytoskeletal genes differentially expressed in cluster 7 compared to its origin, cluster 2. Orange: higher expression in cluster 7, blue: lower expression in cluster 7.

**Supplementary Fig. 7.**
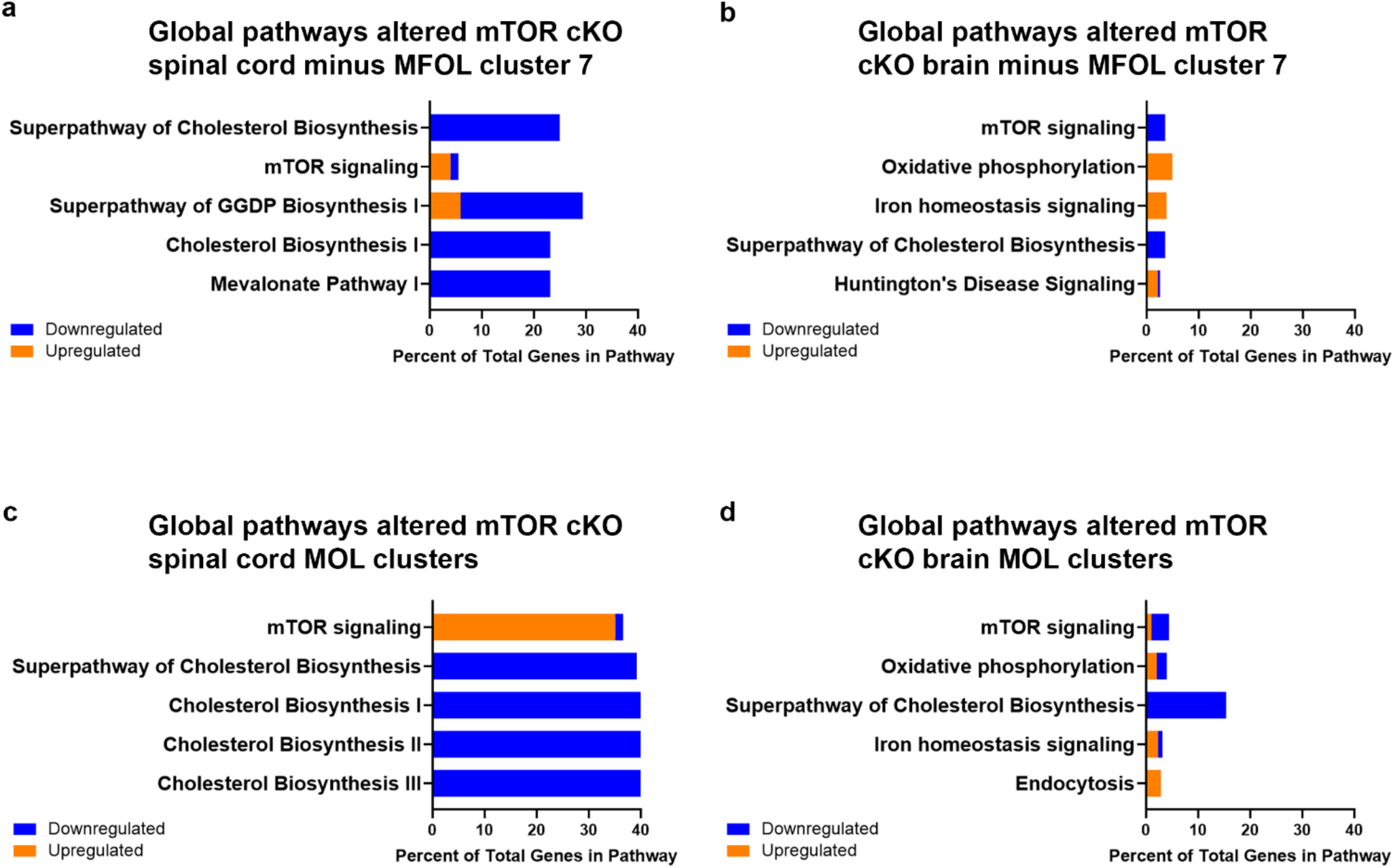
**a, b,**Cells from MFOL cluster 7 were excluded from differential gene analysis. Top 5 pathways dysregulated in mTOR cKO spinal cord (a) and brain (b). **c, d,**Top 5 pathways dysregulated in mTOR cKO MOLs in spinal cord (c) and brain (d). Clusters 1 and 6 were combined for analysis. IPA analysis of genes differentially expressed in mTOR cKO cells compared to control cells; most significantly altered pathways shown. Blue: downregulated genes; orange: upregulated genes.

**Supplementary Fig. 8.**
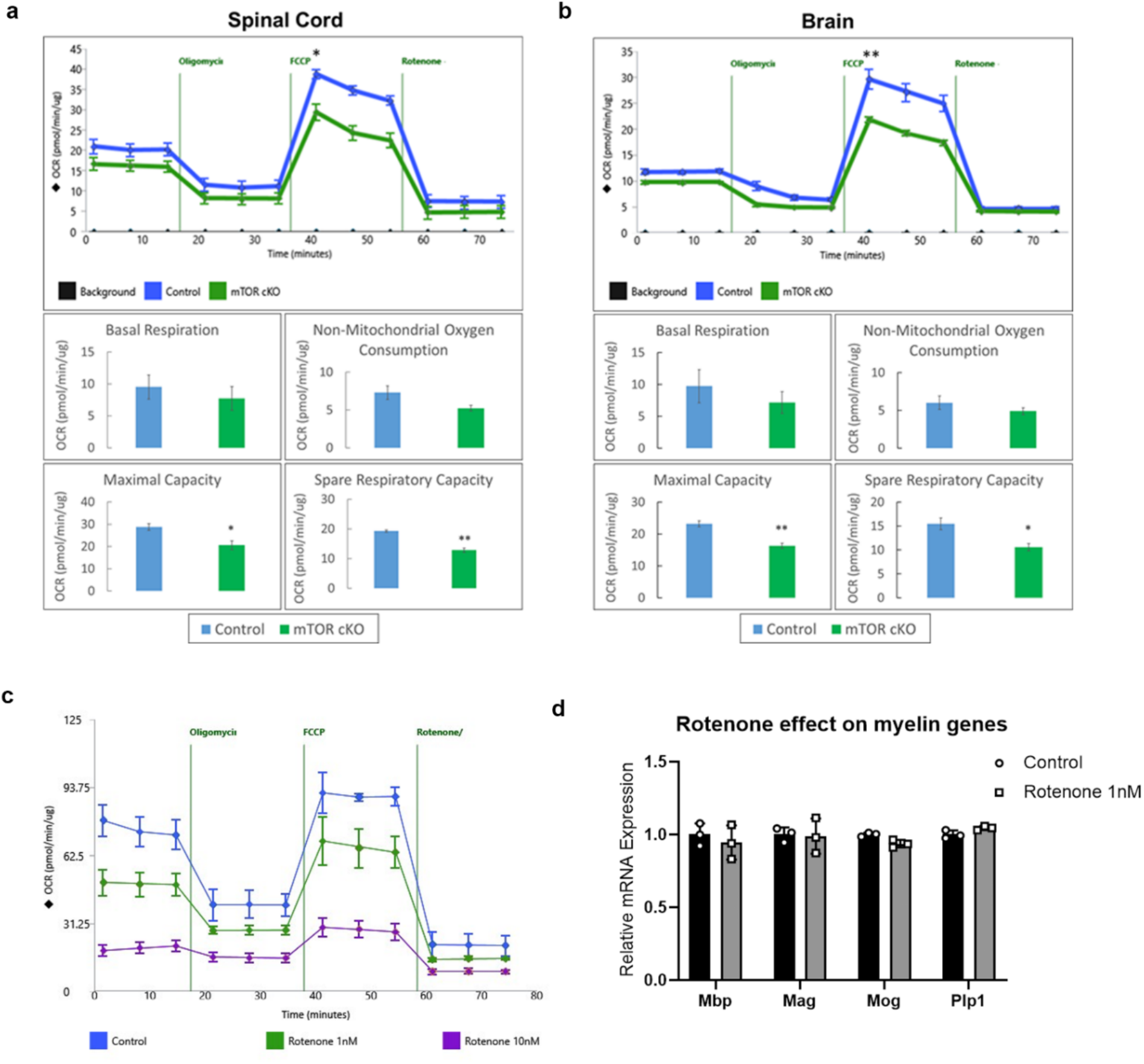
mTOR is necessary for normal oxidative phosphorylation in oligodendroglia. **a-b**, Loss of mTOR reduces maximal respiration and spare respiratory capacity in both spinal cord (**a**) and brain (**b**). O4+ cells were isolated from control and mTOR cKO mice at P10 (spinal cord) and P14 (brain) and plated into Seahorse XFp wells. Oxygen consumption rate (OCR) was measured on Seahorse analyzer using the mitochondrial stress test. Initial measurements were performed to assess basal respiration and again after the addition of the ATP synthase inhibitor oligomycin (0.5 μM). The uncoupling agent FCCP (2 μM) was then added to measure maximum respiration capacity. Rotenone, a complex I inhibitor (0.5 μM) and antimycin A (0.5 μM), a complex III inhibitor, were then added to measure non-mitochondrial oxygen consumption. All Seahorse values were normalized based on total protein content measured after completion. 3 animals/genotype were pooled for each experiment, and experiment was performed 3 times. Values expressed as mean ± SEM, *p ≤ 0.05 **p ≤ 0.01. **c**, Rotenone treatment to reduce OCR to similar levels as loss of mTOR. Differentiating primary rat OPCs were treated with rotenone at 1nM or 10nM for 3 days and then analyzed by Seahorse mitochondrial stress test. Cells were plated and treated in triplicate. **d**, Differentiating primary rat OPCs were treated with 1nM rotenone for 3 days to inhibit OCR to similar levels as loss of mTOR. Expression of *Mbp, Mag, Plp1,* and *Mog* were measured by quantitative real-time PCR. Expression normalized to *Gapdh* as housekeeping gene, then normalized to control. Cells were plated and treated in triplicate. Values expressed as mean ± SD.

**Supplementary Fig. 9.**
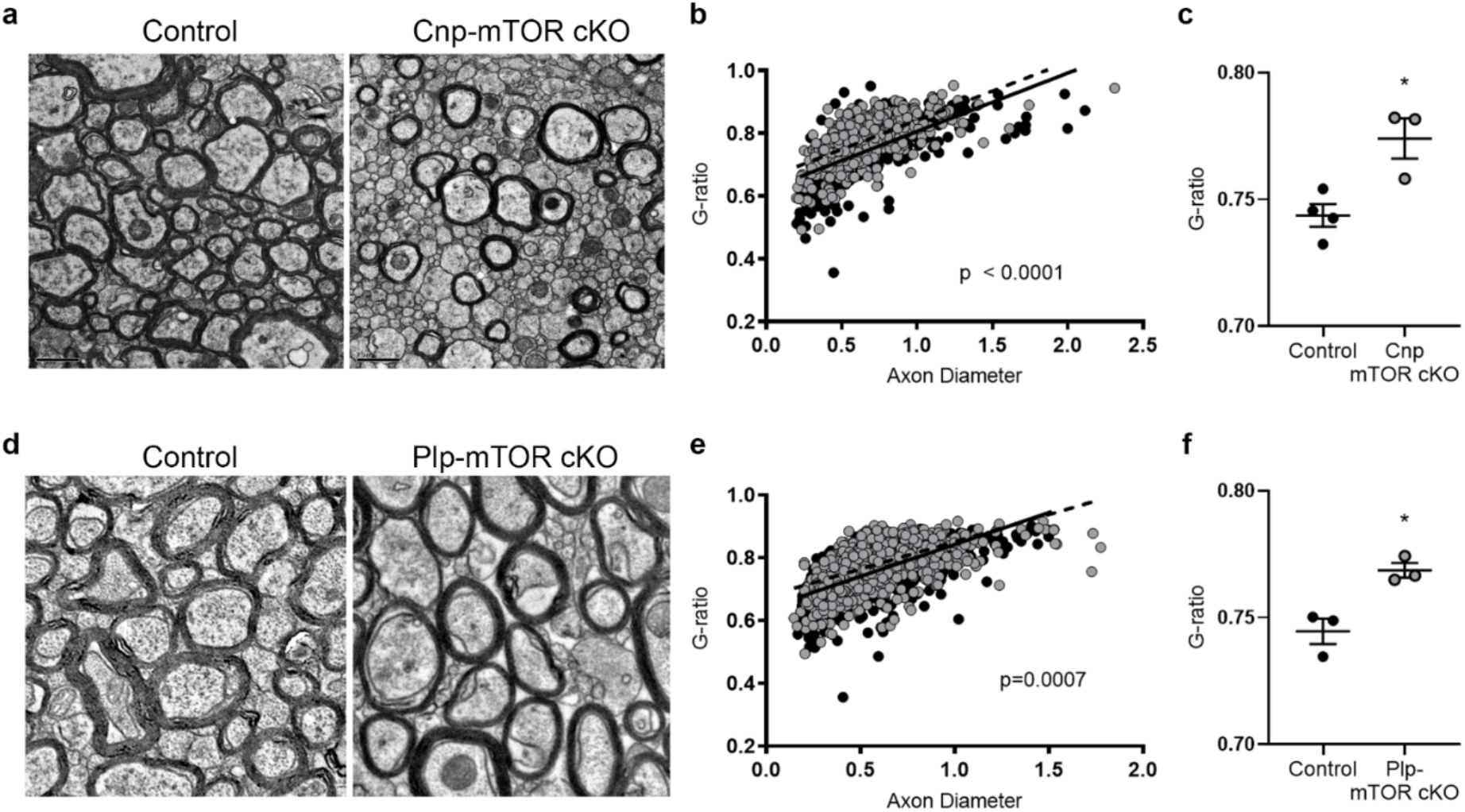
**a**, Representative EM images from 12 months of age control and Cnp-mTOR cKO callosa, scale bar = 1 μm. **b**, G-ratio scatter plot showing linear regressions for control (solid) and Cnp-mTOR cKO (dashed) g-ratios, n > 1400/group. **c**, Average g-ratio of control and Cnp-mTOR cKO at 12 months of age. **d**, Representative EM images from 12 months post-tamoxifen control and Plp-mTOR cKO callosa, scale bar = 1 μm. **e**, G-ratio scatter plot showing linear regressions for control (solid) and Plp-mTOR cKO (dashed) g-ratios, n > 1400/group. **f**, Average g-ratio of control and Plp-mTOR cKO at 12 months post-tamoxifen. For **d-f**, homozygous *mTOR^fl/fl^* mice were bred to heterozygous *Plp-Cre^ERT^*(B6.Cg-Tg(Plp1-cre/ERT)3Pop/J, The Jackson Laboratory, RRID:IMSR_JAX:005975) mice to establish a tamoxifen-inducible *Plp-Cre^ERT^;mTOR^fl/fl^* (Plp-mTOR cKO) mouse line with Cre+/- conditional knock-out mice and Cre-/- control littermates. In the Plp-mTOR cKO mouse CNS, PLP-expressing mature OLs exhibit deletion of mTOR and expression of a membrane-bound GFP reporter. In order to induce recombination, tamoxifen (75 mg/kg; 10 mg/mL stock in 9:1 sunflower seed oil:ethanol) was administered by intraperitoneal injection to adult (8-10 weeks) littermate mice for 5 consecutive days, resulting in recombination of PLP+ mature OLs in adult Plp-mTOR cKO mice. Only male mice were used to avoid endogenous estrogen-mediated Cre leakiness.

**Supplementary Fig. 10.**
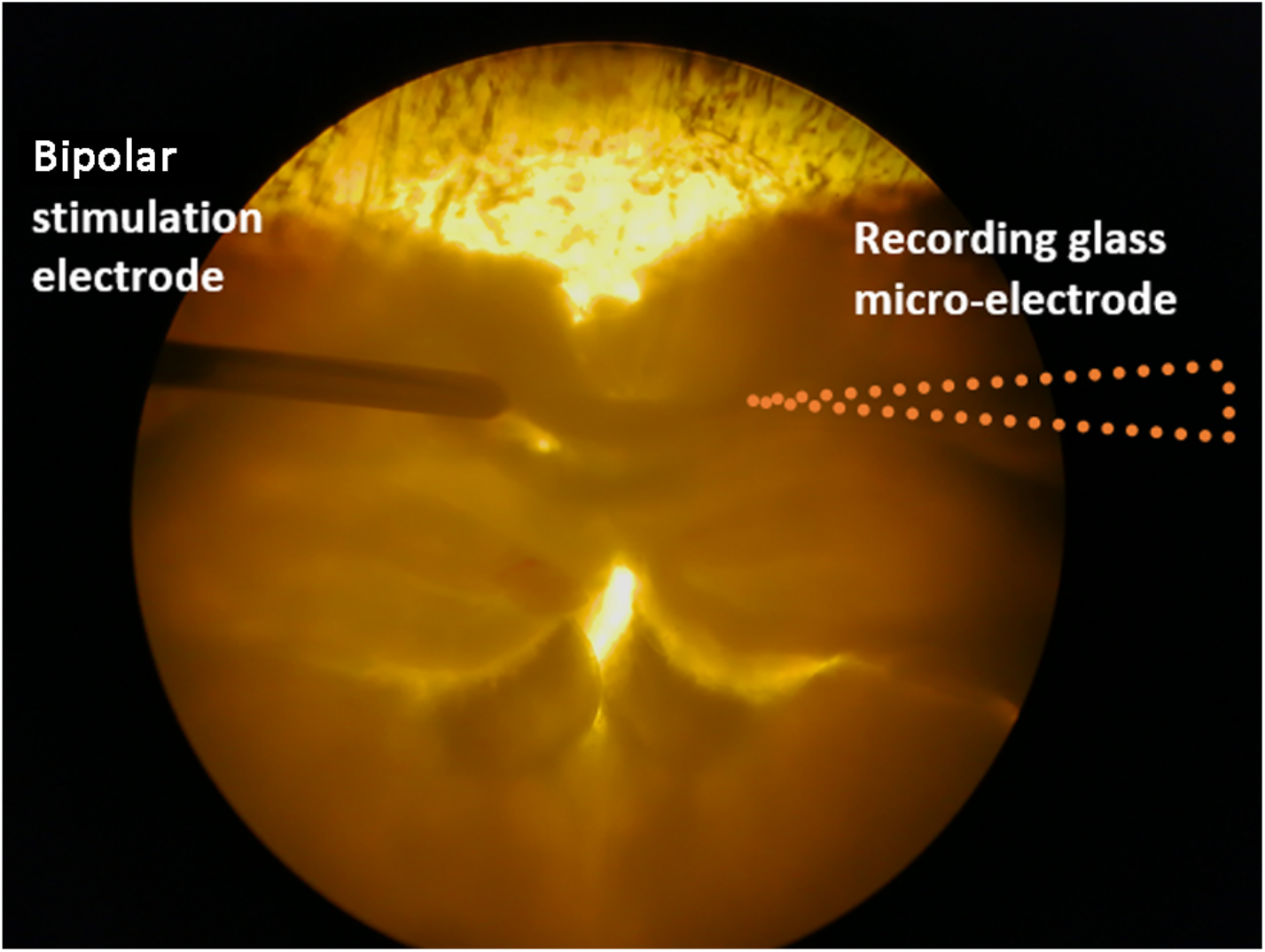
Placement of stimulating and recording electrodes for callosal slice recordings.

**Supplementary Fig. 11.**
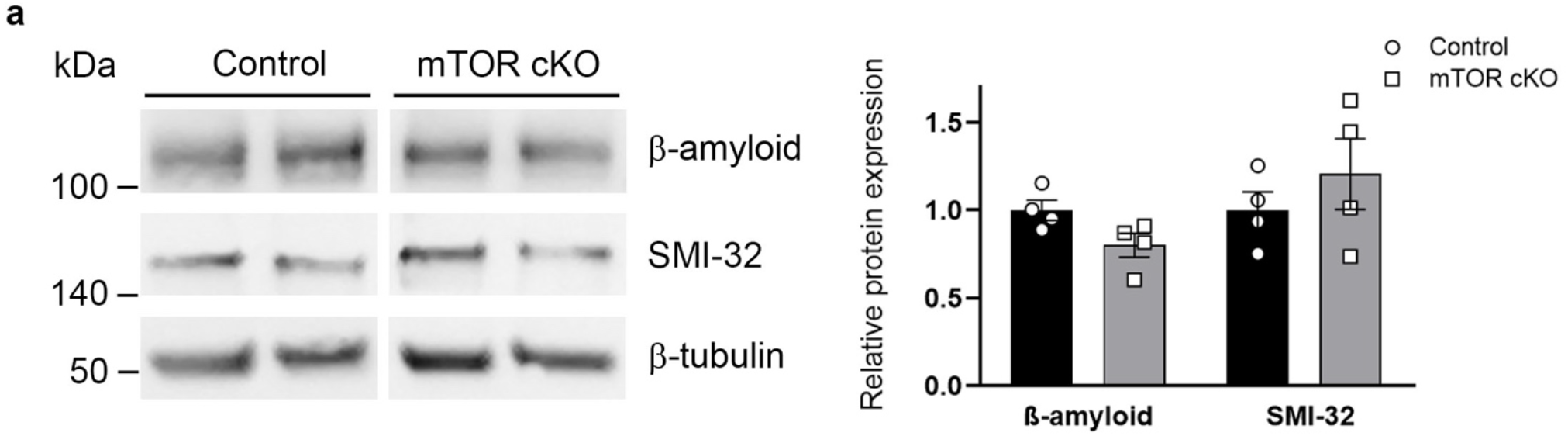
**a**, β-amyloid, an early marker of axonal pathology, and SMI-32, which detects non-phosphorylated neurofilament, expression in 12-week old control and mTOR cKO microdissected corpus callosum. All lanes normalized to β-tubulin, then control. Control and mTOR cKO were run on the same western blot. Graph presents quantification of β-amyloid and SMI-32 from 4 animals/genotype.

